# Context-Dependent Variability Of HIF Heterodimers Influences Interactions With Macromolecular And Small Molecule Partners

**DOI:** 10.1101/2025.05.29.656908

**Authors:** Joseph D. Closson, Xingjian Xu, Meiling Zhang, Tarsisius T. Tiyani, Leandro Pimentel Marcelino, Eta A. Isiorho, Jason S. Nagati, Joseph A. Garcia, Kevin H. Gardner

## Abstract

Hypoxia inducible factors (HIFs) are transcription factors that coordinate cellular responses to low oxygen levels, functioning as an α/β heterodimer which binds a short hypoxia response element (HRE) DNA sequence. Prior studies suggest HIF/HRE complexes are augmented by the binding of additional factors nearby, but those interactions are not well understood. Here, we integrated structural and biochemical approaches to investigate several functionally relevant HIF assemblies with other protein, small molecule, and DNA partners. First, we used cryo-electron microscopy (cryo-EM) to establish HIF-1 and HIF-2 self-assemble to form novel “dimer-of-heterodimers” (DoHD) complexes on extended human EPO enhancer sequences, with one heterodimer bound at a canonical HRE site and the second binding in an inverted fashion to an HRE-adjacent sequence (HAS) 8 bp away. Consistent with ARNT PAS-B domains predominating interactions within a DoHD, we found HIF-1 and HIF-2 co-assemble mixed DoHD complexes on the same DNA. Second, we saw that despite the increased complexities of the larger complexes, ligands for the isolated ARNT or HIF-2α PAS-B domains are still capable of binding and disrupting both the heterodimer and DoHD complexes, albeit with variable potencies depending on the ligand. Finally, we combined cryo-EM and hydrogen-deuterium exchange by mass spectrometry (HDX-MS) to show how HIF-1 and HIF-2 heterodimers engage the transforming acidic coiled-coil containing protein 3 (TACC3) coactivator via both ARNT and HIF-α subunits, though this was unseen in the larger DoHD. Our findings highlight the importance of both molecular context and dynamics in biomolecular complex formation, adding to the complexities of potential regulation.

**Significance Statement:** Hypoxia inducible factors (HIFs) are transcription factors that regulate oxygen-dependent cellular processes with implications in certain types of cancers. Current molecular structures of HIFs bound to short DNA fragments provide insights into their function, but leave open questions about how they bind longer natural DNA fragments and interact with small molecules and protein coactivators. Integrating structural and biochemical techniques, we discovered a novel assembly in which two HIFs bind together on a single extended DNA fragment, forming a “dimer-of-heterodimers”, which exhibits some differences in ligand and coactivator binding than heterodimers or isolated PAS domains. Our studies highlight how functional contexts can shift structural paradigms and provide greater insight into the mechanisms by which HIFs and similar bHLH-PAS transcription factors operate.

## Background

The hypoxia inducible factors (HIFs) are basic helix-loop-helix Per-ARNT-Sim (bHLH-PAS) transcription factors that regulate oxygen dependent processes for higher eukaryotes under hypoxic conditions, controlling the expression of hundreds of genes in erythropoiesis, cell growth, metabolism, and other pathways involved in adaptation to low oxygen levels (3, 11).

HIFs are heterodimers comprised of one of three isoforms of an oxygen-sensitive α-subunit (HIF-1, 2, 3α) and a constitutively expressed β-subunit, aryl hydrocarbon nuclear translocator (ARNT, also known as HIF-1β) (3, 12). Under normoxia, HIF-α subunits are degraded and inactivated through O_2_-dependent enzymatic hydroxylation of specific proline and asparagine residues in the HIF-α C-terminus (4–6). These hydroxylation markers recruit the Von Hippel-Lindau (VHL) E3 ubiquitin ligase (proline hydroxylation) and block recruitment of p300/CREB-binding protein (CBP) transcriptional coactivators (asparagine hydroxylation), coordinately repressing HIF activity via independent mechanisms (3). These hydroxylation events cannot occur under hypoxia, allowing HIF-α to accumulate, enter the nucleus, and heterodimerize with ARNT, resulting in a HIF-α/ARNT complex (HIF complex, HIF) bound to a 5 bp hypoxia response element (HRE) box (5’-RCGTG) in the promoter or enhancer regions of oxygen-regulated genes to regulate transcription (**Fig. 1A**) (3, 11, 13, 14). DNA-bound HIF complexes recruit other regulatory components such as p300 or CBP coactivators, as well as a class of coiled-coil coactivators (CCCs), as integral parts of transcriptional regulation (**Fig. 1**) (10, 11, 15).

**Figure 1.**
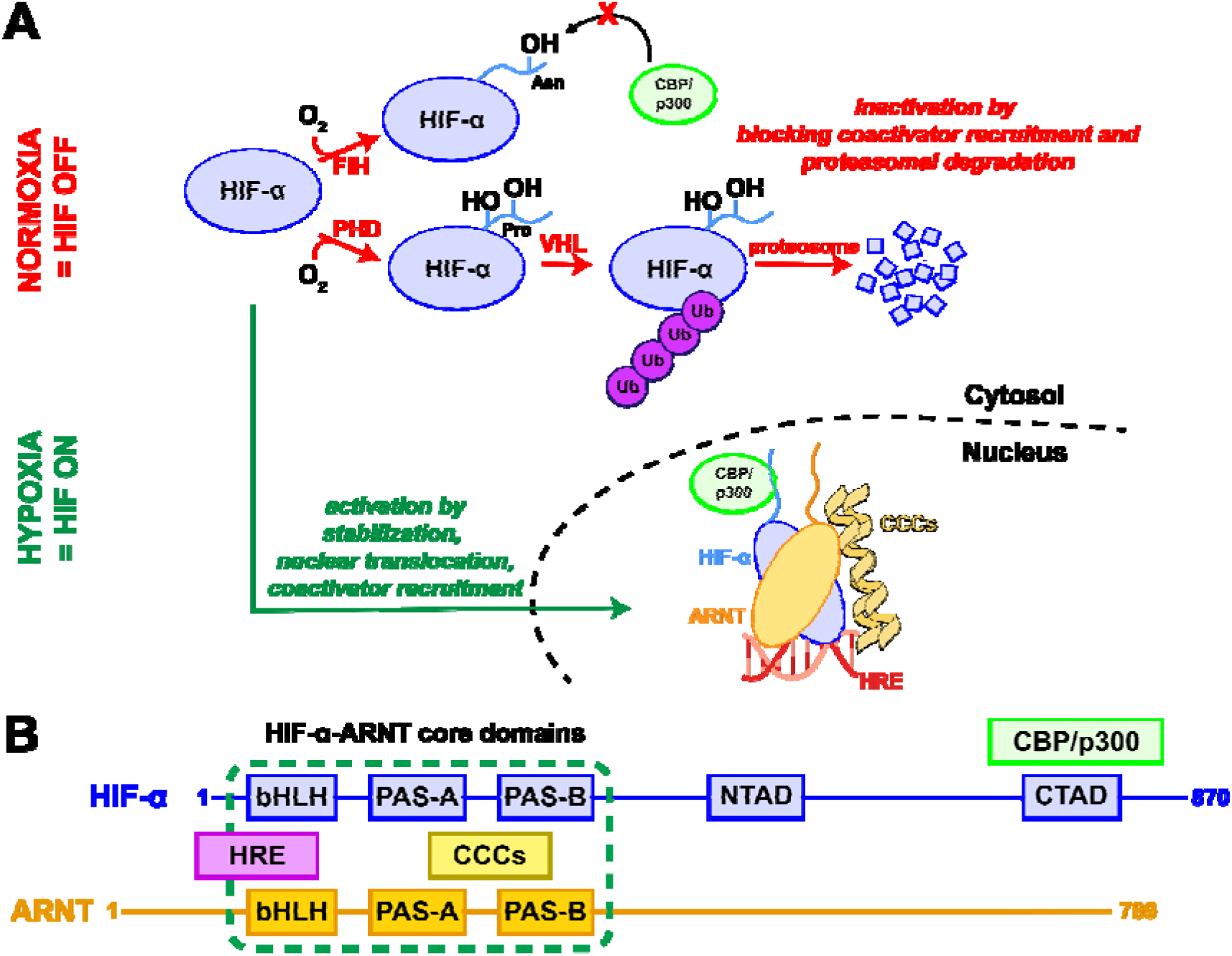
Schematic of HIF pathway regulation and components. (A) Under normoxic conditions, HIF-α is inactivated in the cytosol through parallel oxygen-dependent post-translational hydroxylations of residues in the protein C-terminus. These include a proline-directed mechanism (HIF-1α: Pro402 and 562, HIF-2α: Pro405 and 531) which leads to the recruitment of the von Hippel-Lindau E3 ubiquitin ligase, leading to poly-ubiquitination and proteosomal degradation (3, 4). A second route focuses on an asparagine residue (HIF-1α: Asn803, HIF-2α: Asn851) in the C-terminal transactivation domain (CTAD), preventing recruitment of CBP/p300 coactivators essential for HIF-driven transcriptional activation (5, 6). Under hypoxic conditions, neither of these HIF-α hydroxylations occur, allowing the fully functional protein to accumulate, translocate into to the nucleus, and heterodimerize with ARNT. The HIF-α-ARNT complex then associates with specific HRE sequences in the promoter or enhancer regions of oxygen-dependent genes and recruits coactivators, leading to transcriptional activation. (B) Schematic depicting the domain arrangements of human HIF-2α and ARNT, as well as indicating which domains interact with HRE DNA as well as CCC- and CBP/p300-type transcriptional coactivators.

Functionally, HIF-1 and 2 play major roles in natural cellular hypoxic responses, with their dysfunction implicated in a range of cancers and other diseases (3, 11, 16–18). Though their expression profiles tend to overlap, HIF-1 is typically associated with expression of glycolytic enzymes ubiquitously, whereas HIF-2 is associated with expression of erythropoietin and angiogenic genes localized to the kidney, lung, heart and small intestine (19, 20). Conversely, HIF-3 has been implicated in fatty acid metabolism and potentially as a negative regulator of hypoxic gene expression (21, 22). Constitutively high levels of HIFs can result in increased levels of cellular growth, leading to tumorigenesis and specific types of cancers such as renal cell carcinoma (HIF-2) and breast cancer (HIF-1), while the naturally hypoxic environment of tumors can also upregulate HIF activity, exacerbating tumor progression through activation of hypoxic pathways (11, 16–18, 23). Therefore, HIFs are an attractive target in the development of anticancer agents with one such method of inhibition being the disruption of HIF-α-ARNT complex formation, the proposed mechanism of action for the FDA approved HIF-2 inhibitor and renal cell carcinoma/Von Hippel Lindau syndrome treatment, belzutifan (24–26).

Several protein/DNA crystal structures of bHLH-PAS transcription factors provide structural insights into HIF/DNA binding interactions, most relevantly structures of murine HIF- 1 and HIF-2 bound to artificial 20 bp DNA fragments centered on canonical 5 bp HRE boxes (5’-RCGTG) (9). Collectively, these structures show expected bHLH/DNA interactions, as well as extensive association between the bHLH and PAS domains between both protein subunits (**Fig. S1A,B**) (9). More broadly, this structural arrangement is highly conserved among crystallographic and cryo-EM structures of various HIF complexes, as swapping HIF-1/2/3α isoforms or ARNT homologs (to brain and muscle ARNT-like, BMAL1), leads to only minor perturbations (**Fig. S1**) (9, 27, 28). Substantial changes in organization are only seen with changes in α subunits, as seen between HIF-2α/BMAL1 and CLOCK/BMAL1 (28, 29) complexes, which reorganize quaternary arrangements of PAS domains within the heterodimer assembly (**Fig. S1**).

While these structural studies improved the understanding of protein/protein and protein/DNA interactions in HIF complexes, several fundamental gaps in knowledge remain. Foremost among these is the inability to explain long-standing data showing the importance of DNA sequences adjacent to canonical HREs for HIF-driven transcriptional regulation. Initially identified by deletion/mutation experiments of an enhancer region 3’ to the human EPO gene, full HIF-1 activity depends on the presence not only the HRE-box, but also an additional short sequence (5’-CAC-3’) 8 bp downstream, dubbed the HRE adjacent sequence (HAS) (14, 30). Deletion or mutagenesis of the HAS markedly reduced HIF activity in cells (30); recent follow-up studies showed that even minor changes to the length of the 8 bp HRE/HAS intervening sequence similarly ablates HIF activity in cells, strongly implying cooperativity between these regions (14, 30). While initially suggested that HAS sequences might function by recruiting a separate transcription factor complex such as CREB-1/ATF-1, more recent studies suggest the HAS might bind a second HIF heterodimer due to its partial complementarity to the HRE (14, 31, 32). Such higher order complexes have not been seen in existing structures of HIFs bound to short HRE-only DNA fragments, leaving the arrangements of such higher-order HIF complexes an open topic.

A second open area relates to the binding of small molecules within bHLH-PAS complexes, which can naturally or artificially modulate their function. Most of such compounds bind within the PAS domains themselves, particularly into substantially-sized internal cavities can exist (*e.g.* 100-500 Å^3^ in HIF-α PAS-B domains (27, 33)), providing sites for specific binding to disrupt protein/protein interactions. This strategy has been successfully used for the HIF-2α specific inhibitor belzutifan, now clinically used to treat various forms of clear cell renal cell carcinoma (19, 21, 24–26, 34). Additional binding sites are on PAS domain surfaces (1), providing alternative routes to such regulation. However, almost all binding information on these compounds have been determined on isolated PAS domains (1, 7, 35) or DNA-free bHLH-PAS heterodimers (9, 24, 27). As such, substantive questions remain open about the impact of HIF complex state on small molecule binding.

Finally, information remains sparse regarding interactions between the HIF-α/ARNT heterodimer and other proteins involved in transcriptional regulation, such as transcriptional coactivators. While some of these, such as p300/CBP, are well-known to interact with C-terminal disordered regions in HIF-1α and HIF-2α (36), other coactivators are thought to directly interact with the ordered bHLH/PAS core. These include the CCCs, three of which are known components of ARNT-containing transcriptional complexes: transforming acidic coiled-coil containing protein 3 (TACC3), thyroid hormone receptor/retinoblastoma interacting protein 230 (TRIP230) and coiled-coil coactivator (CoCoA) (2, 15, 37–39). Prior biochemical and structural studies from our group suggested that ARNT/CCC interactions are driven by the ARNT PAS-B domain binding to a C-terminal conserved “TACC box” in TACC3, CoCoA, and TRIP230 (10, 15, 37, 40). However, studies on CCC/ARNT PAS-B complexes (including binding, NMR, and crystallographic analyses (10, 15)) and crystal structures of CCC-free bHLH-PAS heterodimers (9) have conflicting views of the accessibility of proposed binding sites, necessitating characterization of CCC-bound bHLH/PAS complexes to resolve these issues.

With these gaps in knowledge, we assert that current structural models of HIF heterodimers may not fully explain existing data about larger or variant HIF complexes that form in different functional contexts with additional DNA, proteins, or small molecules bound. To address this possibility, we explored different HIF complexes using cryo-electron microscopy (cryo-EM) together with a variety of biochemical and biophysical approaches. Specifically, we compared cryo-EM models of human HIF-2 bound to either a synthetic 20 bp HRE fragment or extended DNAs from the human EPO enhancer region which naturally contain both the HRE and HAS boxes. While we observed a very similar HIF-2/20 bp HRE cryo-EM structure (3.61 Å) as previously seen by X-ray crystallography, the HIF-2/51 bp HRE/HAS cryo-EM structure (3.74 Å) adopted a novel larger configuration, with two individually assembled HIF-2 heterodimers binding each other symmetrically while simultaneously engaging the HRE and HAS boxes of a single DNA fragment. These larger “dimer of heterodimer” (DoHD) complexes are chiefly mediated by ARNT-ARNT interactions between the adjacent HIF-2 heterodimers, implying that they could form in other HIF complexes as well. Consistent with this, we observed longer HRE/HAS DNAs efficiently forming similar large complexes for HIF-1 and even mixed HIF- 1/HIF-2 complexes. Additionally, we investigated the ability of both HIF-2 complexes to bind small molecules known to bind isolated HIF-2α and ARNT PAS-B domains, finding these ligands can still bind and disrupt the larger complexes, despite the apparent inaccessibility of their binding sites in static structures. Finally, we propose a binding interface for CCCs on HIF-1 and HIF-2 bound to 20 bp DNA fragments through cryo-EM modeling and independently validated by hydrogen-deuterium exchange mass spectrometry (HDX-MS). This binding, while observed in heterodimers, is seemingly impaired in the larger DoHD complex, again suggesting differences in dynamics among the complexes. Taken together, our findings suggest that the molecular context of HIF complexes can substantially modulate their structures, which in turn impacts their ability to function and associate with other functionally important components.

## Results

### Cryo-EM of HIF-2/HRE complexes show similar structures, but more dynamics, than crystallographically

Initially, we set out to determine a cryo-EM structure of the human HIF-2 bHLH/PAS heterodimer bound to a synthetic 20 bp HRE DNA fragment previously used for crystallographic studies (9), mixed with C-terminal fragments of human TACC3 or CoCoA. Such a structure would allow direct comparisons between these human complexes and prior structures of murine proteins (**Fig. S1**) (9), while also giving insights into CCC binding modes. The resulting datasets showed a high degree of heterogeneity, including a substantial number of particles that appeared to be HIF-2 complexes without bound CCCs. Focusing first on these HIF-2/HRE complexes, we used WARP (41) to pick ∼1,980,000 particles and cryoSPARC (42, 43) to progressively filter out undesired ones through iterative processes of 2D classification, *ab initio* reconstruction, and 3D refinement (**Fig. S3**). We observed a mix of 2D class averages from this analysis, including those clearly corresponding to HIF-2/HRE complexes like the prior crystal structures (**Fig. 2A**). Intriguingly, we also saw classes that exhibited greater conformational dynamics than previously reported, including those that appeared to be DNA-free (with correspondingly disordered bHLH domains) and DNA-bound complexes with a detached or blurred portion similar in size to a PAS domain (**Fig. 2B**). Given the organization of HIF bHLH/PAS heterodimers, we interpret this as a ARNT PAS-B domain which has separated from the rest of the HIF-2α/ARNT heterodimer.

**Figure 2.**
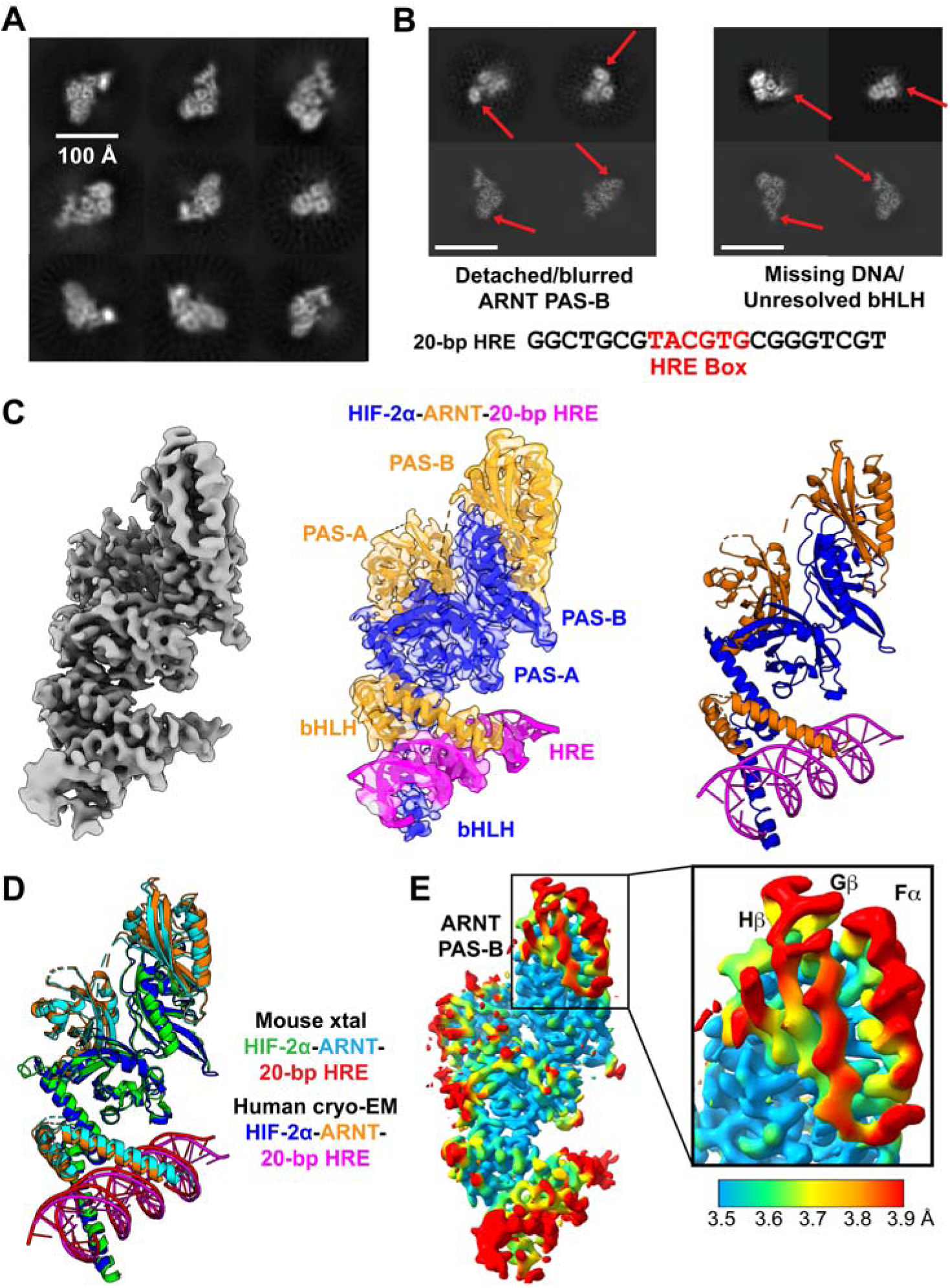
Cryo-EM structural analysis of human HIF-2 bound to 20 bp HRE fragments. (A) Cryo-EM 2D classes of HIF-2 complexes, with individual PAS and bHLH domains and DNA visible. Scale bar represents 100 Å. (B) Some 2D classes of HIF-2 complexes showed either an apparent ARNT PAS-B detached from the rest of the HIF-2α-ARNT core (top row, left) or destabilized bHLH domains lacking bound DNA (top row, right). 2D projections generated through backprojection of the final volume are shown in the bottom row for comparison. (C) 3.61 Å EM map (262,466 particles, 13.3% of all collected particles) was generated from refinement of the most populated *ab initio* volume into which an AlphaFold 2 model of human HIF-2 was fit. (D) Overlay of the mouse HIF-2 crystal structure (PDB code: 4ZPK, (9)) and human HIF-2 cryo-EM structure show highly similar domain architectures. (E) Local resolutions mapped onto the HIF-2 cryo-EM density map show a region of particularly low local resolution/high flexibility in the ARNT PAS-B (Fα-helix, Gβ- strand, Hβ strand) domain and bHLH domains.

Using ∼262,000 particles (13.3% of the total particles picked) from the HIF-2/DNA class noted above, we obtained a 3.61 Å EM map which we successfully fit with an AlphaFold-generated model of human HIF-2α-ARNT bound to 20 bp HRE DNA (**Fig. S4A**) without requiring any large-scale rearrangements (**Fig. 2C**), save for the termini of the HRE fragment which were left unmodeled as a result. Similar to HIF crystal structures (9, 27), several long protein loops were unresolved, with the most notable being the ARNT PAS-A/PAS-B linker (ARNT residues 346-359), likely due to high flexibility in these regions.

Notably, while our cryo-EM structure superimposed well with the murine crystal structure, as expected from the > 95% sequence identities of the HIF-2α and ARNT bHLH/PAS regions, we observed that the structures slightly differ in the position of the ARNT PAS-B domains. This difference seems to be a ∼2.5 Å domain-level shift of the cryo-EM ARNT PAS-B domain away from the adjacent HIF-2α PAS-B domain and the rest of the heterodimer (**Fig. 2D**). Supporting this, local resolution analysis of the cryo-EM map estimated most regions had resolutions between 3.2-4.0 Å, with the lowest protein-associated resolutions corresponding to the ARNT PAS-B domain and in some of the extended loops, again consistent with local dynamics in these regions (**Fig. 2E**). Together with our observation of even larger movements of ARNT PAS-B in the 2D class averages (**Fig. 2B**), we suggest that this domain has markedly more dynamics than previously appreciated, with potential functional links we explore below.

### HIF-2α-ARNT forms a heterotetrameric structure on EPO enhancer DNAs

As noted above, previous studies indicated that additional DNA sequences adjacent to HRE boxes are functionally important for full activation of certain oxygen-regulated gene promoters (14, 30). To explore the structural basis of this observation, we used cryo-EM to examine HIF-2 complexes bound to longer ∼50 bp human EPO enhancer DNA sequences, including both a HRE and HAS box previously demonstrated as necessary for full activation. Using DNAs including both HRE and HAS boxes with additional flanking sequences on either the 5’ (dubbed BS1 HRE/HAS, 51 bp) or 3’ (dubbed BS2 HRE/HAS, 52 bp) ends, we collected cryo-EM datasets of human HIF-2 complexes bound to these longer DNAs, mixed with C-terminal TACC3 fragments. As with our studies of the shorter 20 bp HRE complexes, most particles appeared absent of TACC3, so we focus here on complexes of HIF-2 bound to the 51 bp BS1 HRE/HAS fragments.

Early in our analysis, we observed 2D class averages that were markedly different than seen for HIF-2 bound to 20 bp HRE (**Fig. 2A,3A**). Instead, HIF-2 bound to 51 bp HRE/HAS showed classes with much larger protein density arranged more symmetrically about the DNA, prompting further structural examination of these complexes. Final refinement of data from 164,000 particles (25.0% of all particles picked) (**Fig. S5**), led to the determination of a 3.74 Å structure which neatly fit two individually assembled HIF-2α-ARNT heterodimers as well as the 51 bp BS1 HRE/HAS fragment into a “dimer-of-heterodimers” (DoHD) conformation within our density map without needing any large scale domain rearrangement (**Fig. 3B**). We underscore DoHD complexes were observed by size exclusion chromatography before freezing (**Fig. S6**) and we did not see any comparable higher-order assemblies in our HIF-2/20 bp HRE cryo-EM results, supporting that formation of this larger complex can occur *in vitro*, is dependent on the bound DNA component, and is not an artifact of cryo-EM processing. Conversely, we did not see any smaller assemblies of single heterodimers bound to 51 bp HRE/HAS in cryo-EM, implying the DoHD complex is stable enough to be isolated *in vitro* and remain intact during cryo-EM (**Fig. S6**).

**Figure 3.**
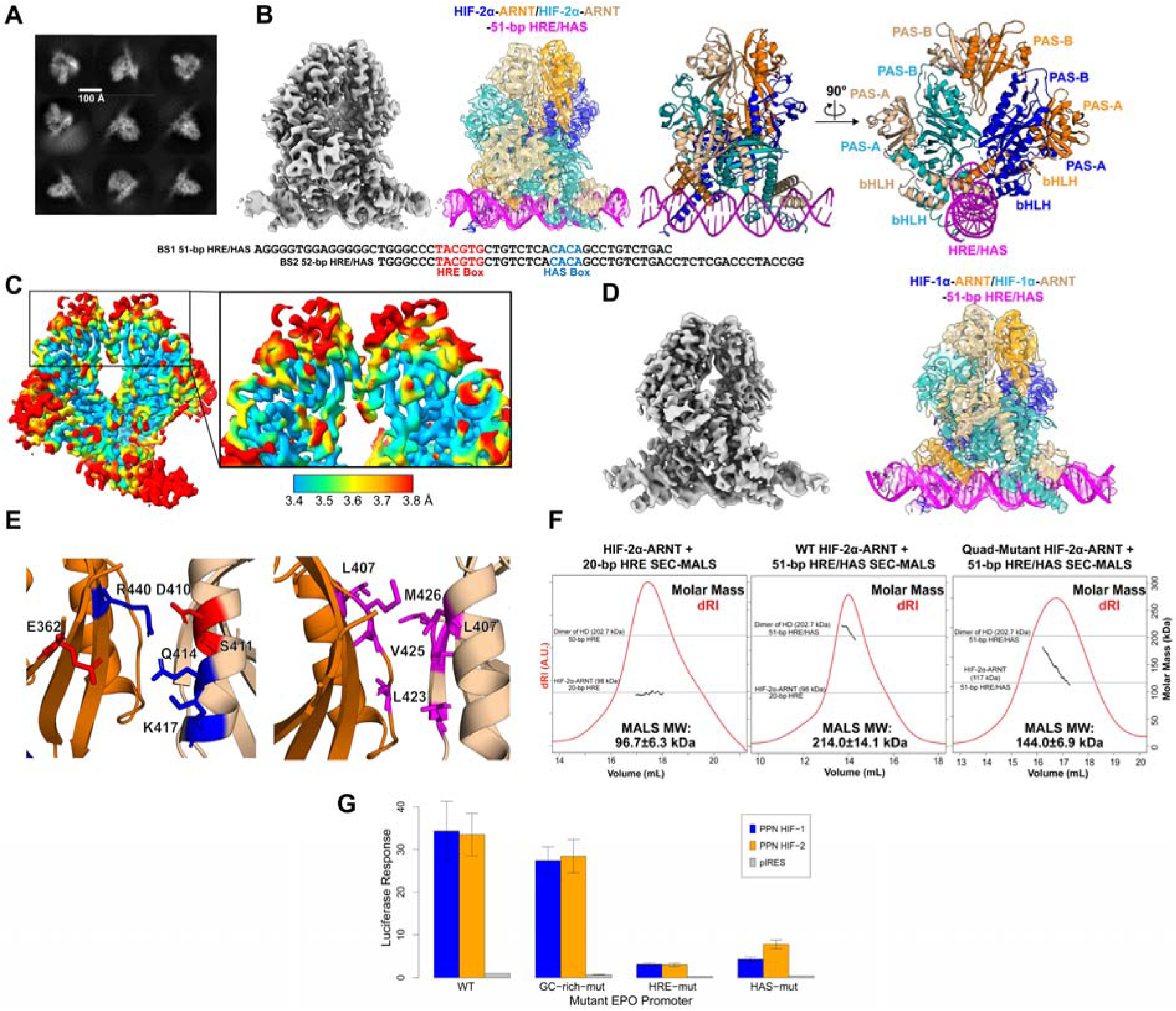
Structural and biochemical analyses of HIF dimer-of-heterodimer complexes. (A) 2D cryo-EM class averages from analyses of data collected on HIF-2α-ARNT heterodimers bound to 51 bp BS1 HRE/HAS DNA fragments, showing markedly different (and often, more symmetrical) arrangements than comparable 2D class averages of HIF-2 bound to 20 bp HRE DNAs. Scale bar represents 100 Å. (B) A 3.74 Å EM map (164,352 particles, 25.0% of all collected particles) was generated from refinement of the most populated *ab initio* volume into which two individually assembled human HIF-2 heterodimers were fit, along with the corresponding 51 bp BS1 HRE/HAS DNA fragment. (C) Local resolution data mapped onto the HIF-2 dimer-of-heterodimers cryo-EM map showed that the ARNT PAS-B Fα-helix and Gβ-strand were higher relative resolution than in corresponding sections of the 20 bp HRE-bound HIF-2 EM map, suggestive of ordering between the heterodimer and dimer-of-heterodimer complexes. (D) The 3.90 Å cryo-EM map (71,802 particles, 27.3% of all collected particles) of HIF-1 bound to a 52 bp BS2 HRE/HAS DNA fragment showed a highly similar domain arrangement to HIF-2, though the ARNT PAS-A domains were less resolved. (E) Highlighted residues were suspected to form interactions between the opposing ARNT PAS-B domains in the HIF-2 supercomplex (basic residues in blue, acidic residues in red, uncharged residues in magenta) (F) SEC-MALS data showed expected differences in the molecular weights of HIF-2 on 20 bp HRE DNA fragments (96.7±6.3 kDa) and 51 bp BS1 HRE/HAS DNA fragments (214.0±14.1 kDa), but revealed an intermediate value (144.0±6.9 kDa) for a HIF-2 variant with mutations in four key ARNT PAS-B Fα-helix and Gβ-strand residues which are at the ARNT/ARNT interface essential to the dimer-of-heterodimers complex. (G) Luciferase reporter assays containing the wildtype (WT) or mutant versions of the human EPO enhancer with changes in the upstream GC-rich region (GC-rich- mut), HRE (HRE-mut), or HAS (HAS-mut) site were performed in HEK293T cells in conjunction with ectopic expression of oxygen-independent (PPN) HIF-1α or HIF-2α. Empty pIRES vector expression was measured to ensure empty vector would not contribute to reporter response. Mutations to either the HRE or HAS site substantially blunted reporter induction by placed at the protein/protein interface despite the limited resolution of the cryo-EM map.

This intriguing higher-order complex orients the two HIF-2 heterodimers around a pseudo two-fold rotation axis, with one heterodimer rotated ∼180° from the other. The first of these heterodimers (1° heterodimer) bound to the HRE box through the HIF-2α and ARNT bHLH domains in a nearly identical fashion to the minimal complexes with 20 bp HRE-only DNA fragments. The 2° heterodimer bound the HAS box through its respective bHLH domains (**Fig. 3B,S6**), positioned in opposition to the 1° heterodimer, with the spacing between the HRE and HAS boxes facilitating direct interactions between the heterodimers. Protein/protein interactions between the two heterodimers were chiefly mediated through the ARNT PAS-B domains, utilizing residues in the Fα-helix and Gβ-strand with potential interactions between Leu407, Gln414, and Met426 of both ARNT PAS-B domains that could be unambiguously ^-^n^2α^ interface despite the limited resolution of the cryo-EM map.

Additional interactions may come from the HIF-2α PAS-A EF loops as well (**Fig. S6**). Cryo-EM data for the central 27 basepairs of the 51 bp fragment were clearly resolved to allow us to confidently place both the HRE and HAS boxes into the cryo-EM map, with slight upward curvature 3’ to the HAS-box before losing resolution at the 3’ terminus, possibly to support binding of the 2° heterodimer to the DNA.

We also performed local resolution analysis of this dimer-of-heterodimers cryo-EM map, which spanned 3.2-4.2 Å resolution (**Fig. 3C**). While similar overall to our analysis from the 20 bp HRE-bound HIF-2, we observed two substantial changes. First, we saw higher resolution density in the dimer-of-heterodimers map corresponding to the ARNT PAS-B domains compared to the 20 bp HRE-bound HIF-2 map, specifically around the β-sheet surface and Fα- helix located at the dimer/dimer interface, implying a domain-level stabilization. In contrast, the dimer-of-heterodimers map appears to show relatively low resolution in the ARNT PAS-A domains of both heterodimers, likely due to being distal to the protein/protein and protein/DNA interactions which stabilize the complex.

### Implications and validation of dimer-of-heterodimer assembly on other HIFs

The central role of ARNT in mediating these dimer-of-heterodimer complexes has several implications. First, analogous HIF-1 DoHD complexes should be able to form on longer DNAs, given the shared role of ARNT in both HIF-1 and HIF-2. Second, point mutations at the ARNT PAS-B domain, which provides the majority of protein/protein interface in the cryo-EM structure, should destabilize dimer-of-heterodimer formation. Finally, “mixed” dimer-of-heterodimers should be evident *in vitro*, with HIF-1 and HIF-2 heterodimers co-complexing to the same DNA fragment. We address each of these implications in turn below.

Starting with the ability of HIF-1 to form dimer-of-heterodimer structures, we collected a cryo-EM dataset of human HIF-1 bHLH/PAS fragments bound to 52 bp BS2 HRE/HASfragments, as initial characterization of HIF-1/DNA complexes appeared more homogenous with BS2 (**Fig. S7**). 2D classes from this dataset were very similar to those from our prior HIF-2/HRE/HAS studies above, suggesting similar structure formation (**Fig. S7B**).

Further data refinement from 71,800 particles led to a 3.90 Å cryo-EM map, into which we could fit two individually assembled HIF-1 AlphaFold-generated models using the inverted repeat-type arrangement as in the HIF-2 dimer-of-heterodimers (**Fig. 3B,D; Fig. S6,S7**). As before, the ARNT PAS-B Fα-helix and Gβ-strand elements appear to provide the bulk of protein/protein interaction surface within the HIF-1 complex, consistent with the ARNT PAS-B domain playing a role in dimerizing HIF-1 as well as HIF-2. The HIF-1α PAS-A EF loops again appear to potentially play a further role in stabilizing the DoHD complex, notably with apparent EM density between nearby cysteines (Cys118) in these loops (**Fig. S7H**). Interestingly, the ARNT PAS-A domains of this complex appeared to be of a much worse resolution than in the HIF-2 DoHD complex, leading to fewer residues of this domain being confidently fit into the EM map (**Fig. S7F**).

### The ARNT PAS-B domain serves as a major interface for HIF-2α-ARNT homodimerization

Our cryo-EM structures and local resolution analyses of the HIF dimer-of-heterodimers support the ARNT PAS-B domain as a dimerization interface between the two HIF-α-ARNT heterodimers (**Fig. 3B-D**). Independently, the ARNT PAS-B domain has been suggested to participate in higher order HIF assemblies via analyses of intermolecular contacts in HIF crystal structures (14), albeit in a different arrangement not supported by our cryo-EM structure. To test the role of ARNT PAS-B as a potential dimerization interface, we mutated specific residues at the ARNT PAS-B/ARNT PAS-B interface of our dimer-of-heterodimer structure. These residues, located in the ARNT PAS-B Fα-helix and neighboring β-strands, are proximal and show improved local resolution in the dimer-of-heterodimers compared to the simpler heterodimer structure, implying these regions are stabilized through direct interactions (**Fig. 3C**). At this interface, we observed several sets of salt-bridges (Glu362 to Gln414/Lys417; Asp410/Ser 411 to Arg440) and nonpolar clusters (Leu423 to Leu423; Met426 to Leu407/Val425) (**Fig. 3E**) which are evolutionarily conserved among vertebrate ARNTs (**Fig. S8**). To disrupt these interactions, we mutated Glu362 and Arg440 to alanine and Leu423 and Met426 to lysine, developing a quad mutant ARNT (ARNT E362A/L423K/M426K/R440A) which we predicted to have lower dimer-of-heterodimer stability than wildtype. Using size exclusion chromatography coupled with multi-angle light scattering (SEC-MALS), we examined the average molecular weights of the wildtype and mutant complexes with the 51 bp BS1 HRE/HAS DNA fragment. For wildtype HIF-2, this complex had an average molecular weight of 214.0±14.1 kDa, similar to the expected 202.7 kDa for two HIF-2 heterodimers bound to a 51 bp DNA duplex and effectively double the 96.7±6.3 kDa observed for a HIF-2 heterodimer bound to a 20 bp HRE fragment (**Fig. 3F**). In contrast, HIF-2 containing the quad E362A/L423K/M426K/R440A mutant ARNT bound to 51 bp BS1 DNA showed a later-eluting SEC peak with an average 144.0±6.9 kDa molecular weight (**Fig. 3F**). Though higher than the expected 117.0 kDa for a single HIF-2 heterodimer stably bound to a 51 bp DNA duplex, our SEC and MALS data strongly support an equilibrium shift towards a single heterodimer, implying disruption of DoHD formation. Combining these results to the cryo-EM structure and local resolution analysis suggest the importance of the ARNT PAS-B Fα-helix and adjacent β- strands as an interface in the formation of dimer-of-heterodimers.

### ARNT-mediated formation of mixed HIF-1/HIF-2 dimers-of-heterodimers

As the last of the three implications of our dimer-of-heterodimer structures, we asked if the commonality of ARNT in mediating comparable HIF-1 and HIF-2 dimers-of-heterodimers could facilitate HIF-1 and HIF-2 complexing together to form a “heterodimer-of-heterodimers” complex on extended DNAs. To test this, we created a fusion protein of HIF-1α tagged on the N-terminus with *E. coli* maltose binding protein (MBP) to increase its molecular weight by about 45 kDa, allowing us to distinguish HIF-1 and HIF-2 complexes by SDS-PAGE and size exclusion chromatography.

After copurifying this MBP-tagged HIF-1 with untagged HIF-2, we bound them to 51 bp BS1 HRE/HAS DNA fragments, and ran this mixed sample over SEC. We observed two peaks in the resulting chromatogram (**Fig. S9A**), with SDS-PAGE analysis showing all three subunits (peak 1, MBP-HIF-1α, HIF-2α, ARNT) in the first, earlier-eluting peak and only HIF-2α and ARNT in the later peak (peak 2) (**Fig. S9D**). The elution volume of the later peak (peak 2) was similar to that of independently isolated HIF-2 DoHD complex while the earlier eluting peak (peak 1) corresponded to an elution volume between those of independently isolated MBP-HIF-1 DoHD and HIF-2 DoHD complexes (**Fig.S9A**). This supports the formation of a tandem HIF- 1/HIF-2 *heterodimer*-of-heterodimers on an extended DNA fragment. Intriguingly, we did not see a peak corresponding to MBP-HIF-1/MBP-HIF-1 heterodimers in the mixed sample, which we would expect to form based on our cryo-EM data and have confirmed through SEC of MBP- HIF-1 heterodimers bound to 51 bp HRE/HAS DNA fragments (**Fig. S9A**).

To explore the DNA sequence requirements for forming these higher order structures, we repeated these experiments using 51 bp BS1 DNA fragments with mutations into either the HRE or HAS-box to reduce protein binding to those sites (**Fig. S9B**). As expected, mutation of the HRE-box prevented almost any heterodimer formation, suggesting the HAS box alone is insufficient to stably recruit either single heterodimers or dimer-of-heterodimers; protein that is not bound to DNA cannot stably form heterodimers, and therefore elutes in the void (**Fig.**

**S9C,S9E**). In contrast, mutation of the HAS-box increased the presence of a later elution peak we attribute to a single heterodimer on DNA, while DoHD formation was reduced but still evident (**Fig. S9C,S9E**).

Coupled with the inability to observe stable dimers-of-heterodimers on 20 bp HRE-only DNAs (by either cryo-EM or X-ray crystallography), we interpret these data as giving us insights into the coupled protein/protein and protein/DNA interactions needed to assemble higher-order HIF complexes: HIF/HRE interactions are key for 1° heterodimer recruitment, with additional DNA interactions – ideally from an appropriately-spaced HAS box – needed to stabilize the recruitment of a 2° heterodimer. This is consistent with prior experiments which scanned the transcriptional potency of HIF-driven promoters with HRE/HAS spacings of 2 to 12 bp (in 2 bp steps) showing maximal activity specifically at 8 bp(14).

### Formation of HIF dimer-of-heterodimers is important to cellular function

Building on our observation of HIF dimers-of-heterodimers *in vitro*, we conducted further studies to determine if these complexes would assemble in cells, and if they are functionally relevant. We generated luciferase reporter plasmids driven by EPO enhancer region sequences or variants containing mutations in the HRE, HAS or 5’ GC-rich regions, and cotransfected these into HEK293T cells together with expression plasmids for oxygen-invariant “PPN” variants of human HIF-1α or HIF-2α. Examining luciferase expression from these cells, we observed robust expression of the reporter genes driven by either the WT promoter or one containing point mutations in the GC-rich region. In contrast, expression from plasmids with HRE mutations was ablated by approximately 90%, as expected from the key role that HREs play in recruiting HIFs. Notably, expression from HAS-mutated promoters was also substantially decreased compared to WT controls, with approximately 70-80% decreased activity (**Fig. 3G**).

As neither mutation completely ablated HIF-dependent transcription of the reporter, we suggest that either the HRE or HAS sequence on its own may still recruit a single heterodimeric complex. Indeed, many HIF-regulated genes do not contain an HAS-box, and therefore must function by recruiting HIF only to the HRE-box (14, 44).

### Structural differences between HIF-2α-ARNT complexes impacts ligand binding

Our work demonstrates that differences in protein and DNA fragments of HIF complexes can markedly impact the quaternary structures of these components. To explore the functional impact of these changes on the binding of small molecules to various HIF complexes, we measured the efficacies of several HIF-2α or ARNT PAS-B binding compounds to disrupt various HIF-2 complexes. Such compounds, which have been sought as leads for disruptors of protein/protein interactions in HIFs, have typically been identified using biophysical or biochemical approaches focused on ligand binding to isolated PAS-B domains (2, 7, 35). These compounds range in affinity from nanomolar (e.g. compound **2** binding to HIF-2α PAS-B, K_d_ 81 nM (7)) to high micromolar (e.g. ARNT PAS-B binding KG-548 and KG-279 (1)) (**Fig. 4A**), but comparable measurements have typically not been available on larger complexes, some of which occlude obvious access to binding sites on or inside of the PAS domain targets. To address this shortcoming, we used amplified luminescent proximity homogenous assay screen (AlphaScreen, Revvity) to measure the ability of several HIF-2α or ARNT targeting ligands to disrupt the HIF-2 heterodimer and the dimer-of-heterodimers (FLAG-tagged ARNT subunits) bound to biotinylated 20 bp HRE or 51 bp BS1 HRE/HAS DNA fragments. AlphaScreen detects the proximity between two biomolecules separately tagged for affinity to either a donor (streptavidin-coated bead for biotinylated HRE or HRE/HAS) or an acceptor bead (anti-FLAG coated bead for FLAG-tagged ARNT) by emitting luminescence in response to diffusion of singlet oxygen between beads. As such, disruption of the HIF-2 complexes would result in displacement of ARNT from the DNA, leading to a detectable decrease in luminescence that can be used to determine the potency of small molecules to disrupt either complex (45, 46).

**Figure 4.**
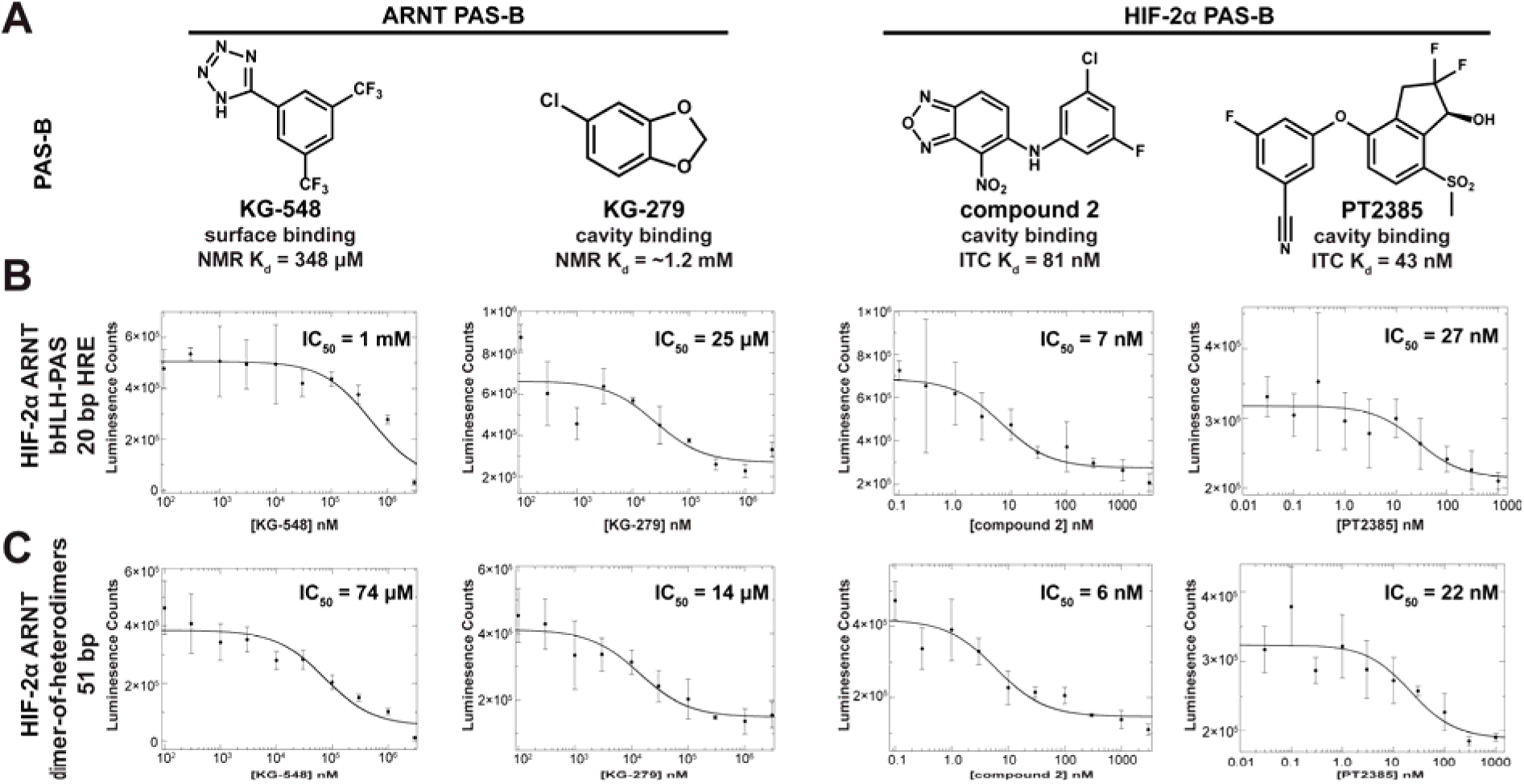
Contextual differences can impact HIF-2 small molecule binding and disruption potency. (A) Known small molecule binders of isolated ARNT (KG-548 (1, 2), KG-279 (1, 2)) and HIF-2α PAS-B (compound 2 (7), PT2385 (8)) domains are shown along with their literature affinities for the specific PAS-B targets, obtained by ITC and NMR. (B) Small molecule disruption of the HIF-2 heterodimer is seen across PAS-B binding compounds, as assayed by AlphaScreen probing interactions between a FLAG-tagged ARNT subunit of HIF-2 and a biotinylated 20 bp HRE fragment. (C) Small molecule disruption of the HIF-2 DoHD complex is also seen across the tested ligands, as detected with AlphaScreen probing interactions between a FLAG-tagged ARNT subunit of HIF-2 and a biotinylated 51 bp HRE/HAS fragment. In both (B) and (C), we observe some differences in disruption potency in the larger contexts compared to the isolated PAS-B domain binding affinity, particularly for the ARNT PAS-B binders.

The first of these compounds, KG-548, disrupted the single HIF-2 heterodimer with a lower potency than the affinity to the isolated ARNT PAS-B(K_d_ 348 µM ARNT PAS-B; ∼ 1 mM IC_50_ HIF-2 heterodimer) despite its binding interface on the ARNT PAS-B surface being blocked by the HIF-2α PAS-B domain in the heterodimeric complex (**Fig. 4B**). This observation suggests some flexibility in the ARNT PAS-B conformation with respect to the rest of the heterodimer, as supported by some cryo-EM 2D classes of the HIF-2 complex with the ARNT PAS-B domain apparently detached from the other core domains (**Fig. 2B**), as well as the lower local resolution of the ARNT PAS-B domain within the 3D heterodimeric complex (**Fig. 2E**). Interestingly, KG- 548 was also capable of disrupting the HIF-2 DoHD complex with a much higher potency (IC_50_ = 74 μM), despite the apparent stabilization of the ARNT PAS-B domains in this complex (**Fig. 4C**). The greatly increased potency of KG-548 against the DoHD complex may suggest new disrupting interfaces for KG-548, possibly between the heterodimers, or a modification of the existing inhibitory interface that increases potency of the inhibitor. We observed KG-279 disrupting both HIF-2 bound to 20 bp HRE DNA fragments (**Fig. 4B**) and HIF-2 DoHD on 51 bp BS1 HRE/HAS DNA fragments (**Fig. 4C**). Though binding to a single heterodimer would be expected due to observed flexibility of the ARNT PAS-B domain, binding to the larger dimer-of-heterodimers suggests the ARNT PAS-B internal cavity is accessible regardless of quaternary structure. The very substantial increase in potency to the HIF-2 heterodimer (IC_50_ = 25 μM) in comparison to the affinity to the isolated ARNT PAS-B (K_d_ = 1.2 mM) suggests novel binding sites for KG-279 on the larger complexes. Additionally, compound 2 and PT2385 were able to disrupt both the HIF-2 heterodimer and DoHD complexes, suggesting the HIF-2α PAS-B cavity remains equally accessible despite the changes in quaternary structure (**Fig. 4C**).

### TACC3 interacts with loops of both HIF-α and ARNT subunits

Though we can assemble dimer-of-heterodimers complexes on HRE/HAS DNA fragments, we understand that only a subset of HIF-driven promoters contain both binding motifs (**Table S1**). Therefore, gene promoters containing only HRE binding motifs, we suspect that HIFs may form varying higher-order complexes with other parts of the transcriptional regulatory machinery. One well studied candidate for HIF binding partners are the CCCs. While CCCs are clearly involved in transcriptional initiation from HIF-driven promoters, and that this is mediated by direct interaction between HIF and CCC proteins, the location of such interaction remains unclear. NMR and other biophysical data from our lab suggested that CCCs bind to isolated ARNT PAS-B domains at a site (2, 15, 39) inconsistent with subsequently-determined crystal (9) and cryo-EM (*vide infra*) structures of larger HIF-2/HRE complexes.

To begin addressing this incompatibility, we sought to characterize larger HIF-2 complexes bound to CCCs. We purified C-terminal fragments of two CCCs, CoCoA and TACC3 (10, 15), confirmed that these were dimeric and coiled coil (**Fig. S10A,B**), and used MST to measure the binding of these fragments to isolated ARNT PAS-B domains. We observed micromolar affinities for these interaction, K_d_ 80 µM for CoCoA and 3 µM for TACC3 (**Fig. 5A**), consistent with prior results (10, 15). Repeating these measurements with the same CCC fragments binding to HRE-bound HIF-2 complexes, however, showed K_d_ ∼10 µM interactions of both coactivators with HIF-2 (**Fig. 5B**). These shifts in affinity between ARNT PAS-B and the intact bHLH/PAS heterodimers show the involvement of additional regions outside of ARNT PAS-B in the HIF-2α/ARNT heterodimer, regardless of whether this is an improvement (CoCoA) or worsening (TACC3).

**Figure 5.**
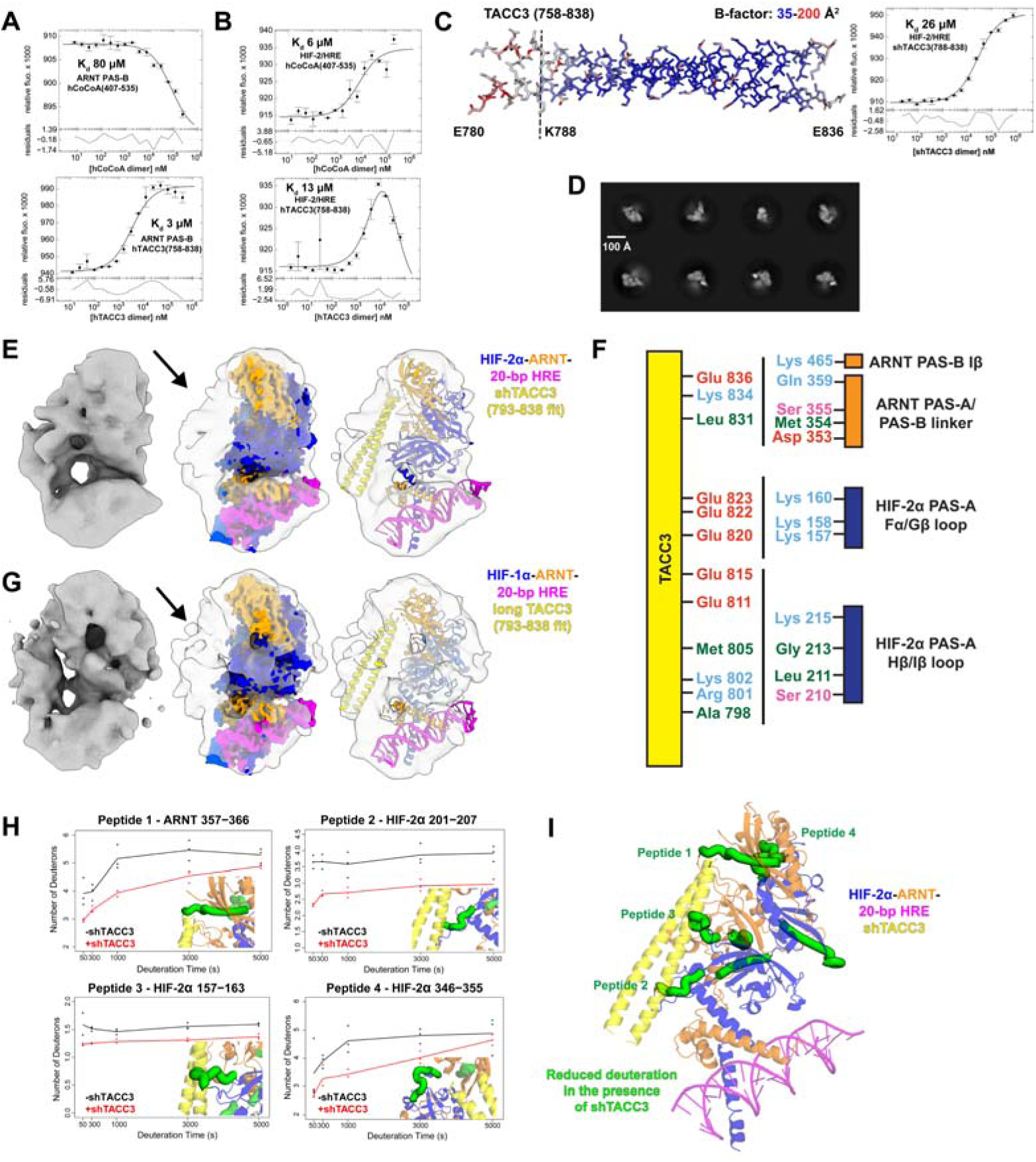
Coiled-coil coactivator binding to HIF heterodimers. (A, B) MST measurements of C-terminal fragments of CoCoA(407-535) and TACC3(758-838) binding to the (A) isolated ARNT PAS-B domains and (B) HIF-2α-ARNT heterodimers (HIF-2) bound to 20 bp HRE fragments. The two highest points of the hCoCoA to HIF-2 binding curve showed evidence of aggregation and were removed. TACC3 binding to HIF-2 showed a decrease at high concentration, suggesting a second binding mode or aggregation. (C) A truncated TACC3(788-838, “shTACC3”) was designed based on our TACC3(758-838) crystal structure showing highly elevated B-factors and poor electron density in residues 758-788. This CCC fragment bound a 20 bp HRE-bound HIF-2 with a minimal affinity drop from the longer TACC3 (Kd 26 μM by MST), and without the secondary binding mode seen for longer fragments to HIF-2. (D) 2D classes of shTACC3 bound to 20 bp HRE-bound HIF-2 showed visible core domains with excess blurred density extending out of the central region. (E) Ab initio reconstruction of 20 bp HRE-bound HIF-2 bound to shTACC3. HIF-2 fit into the volume without major rearrangement, leaving excess density along the ARNT PAS-B/HIF-2α PAS-A interface for most of shTACC3 to be fit. Fitting was solely based on excess density in cryo-EM map of HIF/TACC3 complexes, with the sole assumption that the TACC3 C-terminus is closest to ARNT PAS-B per prior deletion experiments (10). (F) Schematic of interactions between TACC3(788-838) and HIF-2 shows three of the four PAS domains interacting with the TACC3 coiled coil. Residues indicated on TACC3 originate from either chain. (G) Cryo-EM of HIF-1 bound to the longer TACC3(758-858) fragment showed similar excess density as HIF-2/TACC3(788-838), consistent with TACC3 binding similarly to HIF-1 and HIF-2. TACC3 residues 793-838 were fit into the excess density for both HIF-1 and HIF-2. (H) HDX-MS deuterium uptake plots showed several HIF-2α and ARNT loops with increased protection from solvent in the presence of TACC3. (I) Mapping peptides with TACC3-induced HDX changes onto shTACC3-bound HIF-2 model (increased protection in green) showed these protected loops along the proposed TACC3 binding interface from cryo-EM.

To obtain more structural insight into CCC/HIF-2 interactions, we added a TACC3 fragment (residues 758-838) to cryo-EM samples of HIF-2 bound to 20 bp HRE fragments. 2D classes of this complex failed to resolve the HIF-2 protein domains, instead showing blurred circular densities with “stick-like” objects protruding from them, which are of the right dimensions to be TACC3 (**Fig. S10C**). Notably, of these classes showed a bend or break in the stick-like density we attribute to TACC3, suggesting flexibility or dynamics within the coactivator. This hypothesis was supported by a crystal structure of TACC3(758–838) (**Fig. 5C, S10D**), which had low quality electron density and high B-factors for most residues N-terminal of Lys788 (**Fig. 5C, Fig. S10D**), consistent with the break point seen in the cryo-EM classes.

Given this, we made a shorter TACC3(788–838) fragment (“shTACC3”) which we utilized forour subsequent biochemical and structural analyses. MST binding analysis of shTACC3 fragment titrated into HIF-2/HRE complex showed a 26 µM affinity (**Fig. 5C**), slightly weakened from the longer 758-838 construct.

Having confirmed *in vitro* binding on shTACC3 to HIF-2/HRE complexes, we collected a cryo-EM dataset to obtain structural information on this activator/coactivator complex. Using 3,320,000 particles from this dataset, we proceeded with 2D classification and observed classes with visible core domains of HIF-2 as well as blurred excess density which we attributed to shTACC3 (**Fig. 5D**). *Ab initio* reconstruction from 222,000 particles led to a low resolution cryo-EM density map (**Fig. S11**) in which we were able to loosely fit our cryo-EM structure of HIF-2 bound to 20 bp HRE (**Fig. 5E**). This fitting left additional unoccupied density alongside the ARNT PAS-B and HIF-2α PAS-A domains (**Fig. 5E**). Into this region, we fit an AlphaFold structure of the C-terminal 46 residues of shTACC3 as a dimeric coiled coil (**Fig. S5C**) with the C-termini facing the ARNT PAS-B domain, as suggested by our prior work (10, 15). This density became more diffuse towards the shTACC3 N-termini, consistent with the apparent flexibility seen in 2D class averages of longer TACC3/HIF-2 complexes and our TACC3 crystal structure. This structural model places shTACC3 adjacent to the HIF-2 PAS domains, suggesting shTACC3 interacts via three sets of interactions with HIF-2 through unresolved loops in the ARNT PAS-B and HIF-2α PAS-A domains (**Fig. 5F**) and explaining the differences in affinity we observed for TACC3 binding to ARNT PAS-B and larger HIF-2 complexes. Parallel cryo-EM studies of complexes of the longer TACC3 (758–838) bound to HIF-1/HRE showed TACC3- associated density in a comparable spot relative to the HIF heterodimer, suggesting comparable binding interfaces between the two isoforms (**Fig. 5G**). We underscore that while the low resolution of this map implies some inherent motion or flexibility in the HIF and TACC3 orientations, this model explains independently-acquired data from HIF-1/TACC3 and HIF- 2/TACC3 samples as well as MST data suggesting interactions of TACC3 with ARNT PAS-B and additional parts of the HIF heterodimers.

To further bolster our HIF-2/TACC3 binding model, we acquired HDX-MS data on HIF- 2/HRE complexes in the absence and presence of shTACC3. These experiments revealed several HIF-2α and ARNT peptides with notably less deuteration in the presence of shTACC3, most of which were along the TACC3-binding interfaces seen in cryo-EM. Integrating the cryo-EM and HDX-MS results, we found four loops in the proposed CCC binding interface supported by both methods (**Fig. 5H,I; Table S4**): ARNT PAS-A/B linker (357–366), HIF-2α PAS-A Fα/Gβ loop (157–163), HIF-2α PAS-A Hβ/Iβ loop (201–207) and HIF-2α PAS-B C-terminal loop (346–355). We also observed several HIF-2α peptides with TACC3-induced protection from HDX exchange at sites remote from the proposed binding site, suggesting long-range stabilization of the HIF- 2α/ARNT heterodimer upon CCC binding (**Fig. 5I; Table S4**).

As previously mentioned, we did not see TACC3 binding to the HIF-2 dimer-of-heterodimers via cryo-EM, despite the presence of the CCCs within samples used in these analyses. We performed microscale thermophoresis to further probe interactions between TACC3 and the HIF-2 dimer-of-heterodimers complex, but were again unable to see binding, prompting us to believe TACC3 cannot bind the larger complexes. Notably, the HIF-2α and ARNT loops we saw as critical for binding appear to be unobstructed in the HIF-2 dimer-of-heterodimers complex, leading us to suspect that differences in dynamics across the complex (such as those suggested by differences in cryo-EM local resolution) may contribute to this finding.

## Discussion

HIFs have been implicated in specific types of cancer and the progression of tumorigenesis, making them a target for anticancer therapeutics (16, 17, 25, 26). Development of such drugs, including the HIF-2α inhibitor belzutifan, required a mix of approaches applied to reductionist models of HIF-α and ARNT components typically restricted to the isolated PAS domains, though recently advanced to DNA-bound bHLH/PAS heterodimers. While these approaches were successful at informing both basic science and applications of HIF biology, they left unexplained prior findings in three areas – larger DNAs, small molecule binding, and CCC binding – which we advance here.

The formation of higher-ordered HIF complexes was first suggested by Semenza and Wang alongside their discovery of HIF-driven gene expression, and has been recently bolstered by cellular and biochemical work from Rosell-Garcia *et al.* demonstrating that HRE or HAS mutations within HIF-driven promoters drastically reduce HIF activity in cells, similar to our results in **Fig. 3G** (14, 30, 47). Using a split-enzyme reconstitution assay, these authors also nicely demonstrate the likely formation of DoHD complexes in cells. Taken together, HIF-driven transcription clearly occurs from HRE-only or HRE/HAS-containing promoter/enhancer sequences, with selectivity likely stemming from differences in HIF concentrations. Rosell-Garcia *et al.* proposed that larger HIF complexes would form through interactions between ARNT PAS-B and bHLH domains of interacting heterodimers, as suggested through HIF crystal contacts, though such an arrangement places the two sets of bHLH DNA binding domains relatively far from each other, requiring substantial distortion to bind a single DNA containing HRE and HAS boxes (14).

In contrast, our DoHD complex places two HIF heterodimers in a symmetric arrangement via the ARNT PAS-B β-sheet surfaces, with two sets of DNA binding domains ideally placed to contact HRE and HAS boxes with the optimal 8 bp spacer, and not ones with shorter or longer separations as shown by Rosell-Garcia *et al.* (14). Additionally, the sequences of the HRE- (RCGTG) and HAS-boxes (CAC) being imperfect inverse repeats likely explains how HIFs can recognize this second binding site and bind with a 180° rotation relative to the 1° heterodimer.

Notably, little rearrangement of the HIF heterodimer is required to assemble these larger dimer-of-heterodimer assemblies, in stark contrast to CLOCK/BMAL1 in which heterodimers undergo a dramatic 90° intramolecular pivot as they go from isolated heterodimers to a nucleosome-bound complex (**Fig. S12**). Our model also explains mutation data from Rosell-Garcia showing that intact ARNT PAS-B domains are required for higher-order HIF complexes to form *in vitro* or in cells. Further, the high conservation of the ARNT PAS-B domain across vertebrates, including residues involved in DoHD interactions, suggests that such complexes are likely to be found in a wide variety of HIF species variants (**Fig. 8C**). As such, we suggest that our dimer-of-heterodimers model best explains a variety of biochemical and molecular data on interactions and functional importance of these higher-order HIF structures.

Categorization of different HIF-associated promoters containing HRE-only or HRE/HAS motifs reveals that the latter group are largely involved in cell proliferation, metabolism and angiogenesis, while HRE-only containing genes share these functions along with others including transcription regulation, pH regulation, signaling and drug resistance (**Table S1**) (3, 44). HIFs have about a 2-3-fold higher affinity for the HRE-box alone in comparison to the HRE/HAS-boxes together (14), suggesting a dose dependence in which the formation of higher-ordered complexes, and therefore the expression of HRE-only or HRE/HAS-containing genes, depends upon HIF concentration (**Fig. 6**). This is supported by data from transfection-based control of HIF levels for cells grown in culture (14), but has yet to be demonstrated *in vivo* or with natural or artificial regulators of HIF complexes.

**Figure 6.**
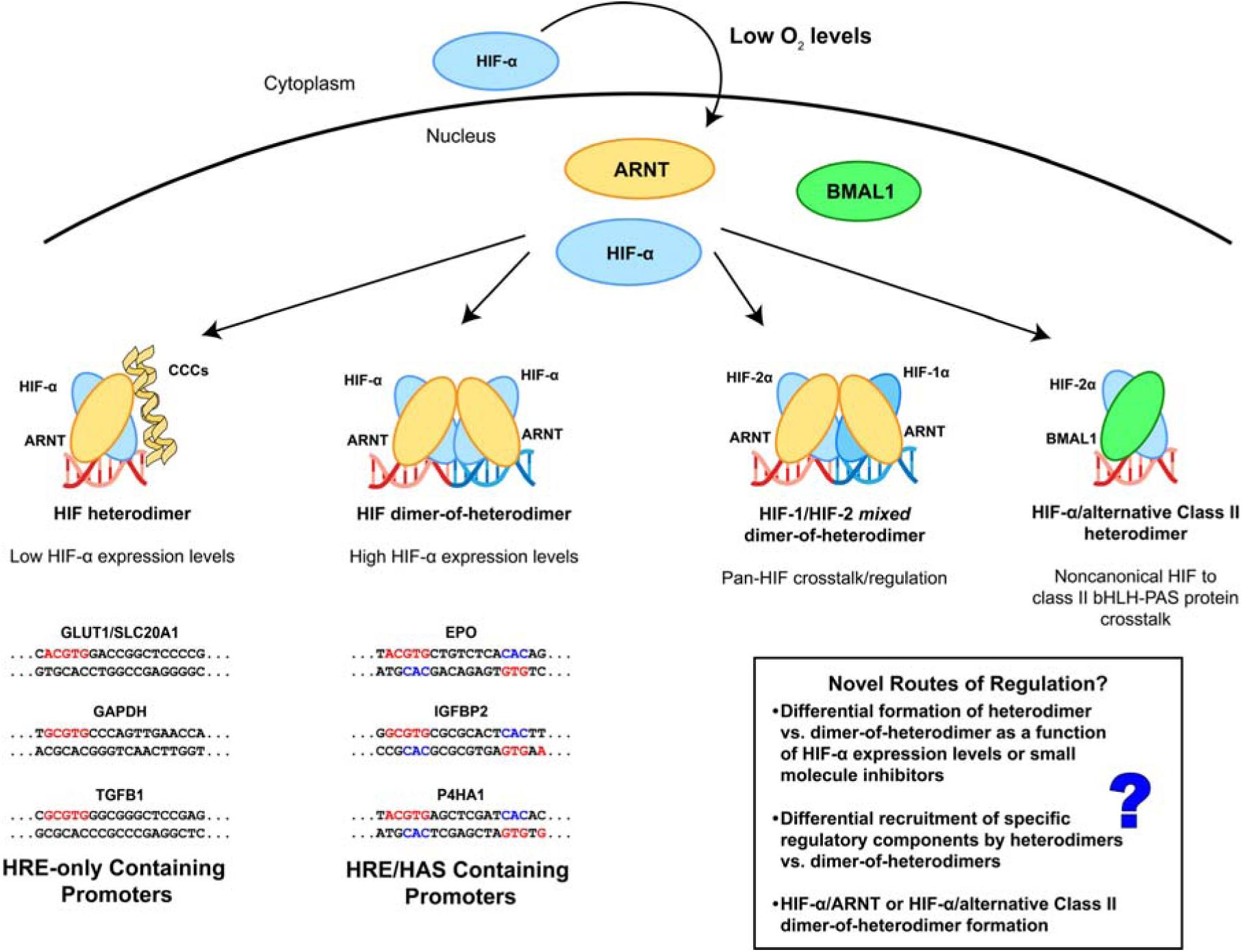
Proposed promoter-dependent formation of higher-order HIF and their downstream effects. (A) Under hypoxic conditions, we propose HIFs form different complexes based on the sequence of the promoter/enhancer region of the gene of interest. For HRE-only containing genes, we propose HIFs form a single heterodimer on the HRE-box, recruiting other components such as CCCs. For genes containing an HRE-box followed by an 8 bp spacer and an HAS-box, we propose that HIFs instead form a dimer-of-heterodimers. We suggest the specification of which genes are activated by HIF are dose dependent, in that lower cellular levels of HIF will activate HRE-only genes while higher cellular levels of HIF will activate HRE/HAS containing genes. Additionally, HIF-α can form noncanonical complexes with alternative class II bHLH-PAS proteins, including BMAL1, as well as mixed HIF-1/HIF-2 dimer-of-heterodimer complexes, each under varying conditions. We suggest that the similar quaternary structure of the HIF-2α-BMAL1 complex lends itself to the potential formation of dimer-of-heterodimers structures, though this remains to be experimentally validated.

The dimer-of-heterodimers concept also strongly suggests the potential for mixed HIF- 1/HIF-2 complexes controlling transcription alongside the more canonical HIF-1/HIF-1 or HIF- 2/HIF-2 complexes. Since we have determined both HIF-1 and HIF-2 can co-complex to form mixed dimer-of-heterodimer complexes together on the same DNA fragment, the two isoforms likely engage in crosstalk to regulate overlapping gene expression. Mixed dimer-of-heterodimer formation may also serve as a mechanism of rescue in cases where either HIF isoform is depleted, either through natural or artificial means (*e.g.* belzutifan inhibition of HIF-2). Rescue of either HIF isoform has been seen in cases where HIF-1 is inactivated, HIF-2 activity will increase to compensate (48, 49); we propose that part of this mechanism may be in the substitution of heterodimers in dimer-of-heterodimer complexes.

This observation raises the potential of other bHLH/PAS heterodimers to join in higher order complexes. Recent cryo-EM studies of a HIF-2α/BMAL1 complex show a similar heterodimeric structure as HIF-2α/ARNT, exposing a similar surface on the BMAL1 PAS-B as used in ARNT PAS-B in the formation of DoHD complexes (50) (**Fig. 6**). Sequence alignment of the human ARNT, BMAL1 and ARNT2 PAS-B domains shows that three residues (E362, Q414, R440) integral to DoHD formation are conserved (**Fig. 3E,F;Fig. S8A,B**). With this, we suggest that dimer-of-heterodimer formation may be plausible for other bHLH-PAS transcription factors provided they adopt a similar quaternary structure to HIF. For example, CLOCK-BMAL1 heterodimers do not appear to be compatible with joining HIF/ARNT heterodimers based on existing structures, although some evidence of CLOCK/BMAL1 higher order complexes have been seen in structural studies of nucleosome-associated structures (28, 51, 52).

Switching from the self-assembly of HIF-2α and ARNT to their ability to bind small molecule ligands, our data clearly indicates that small molecule binding and their abilities to disrupt dimerization is conserved across complex assemblies. We have previously found a slate of small molecules which can bind the isolated HIF-2α or ARNT PAS-B domains (e.g. compound 2 (7), PT2385 (8), KG-548 (1, 2), KG-279 (1, 2)) as assessed by NMR, ITC, or MST. Notably, all small molecule compounds we tested disrupted DNA-bound HIF-2α-ARNT heterodimers. Our examinations of KG-548, a small molecule that binds the beta-sheet surface of ARNT PAS-B, shows that it clearly binds and disrupts single HIF-2α-ARNT heterodimers bound to a short HRE-only DNA, despite the fact that its binding surface on ARNT PAS-B is occluded in both our cryo-EM and prior crystallographic structures of these complexes (1, 9).

We view this result as strongly suggesting the ARNT PAS-B domain having more flexibility than evident in these static structures, as supported by the relatively low local resolution of this domain in the single heterodimer/DNA complex and our observation of 2D classes where the ARNT PAS-B seems to have detached from the result of the heterodimer. More broadly, studies of other bHLH/PAS transcription factors show that the ARNT PAS-B domain (or its analog in BMAL1) adopt a very diverse set of conformations with respect to different Class I heterodimerization partners (HIF-α, AHR, NPAS, CLOCK), supporting this hypothesis (9, 29, 53, 54). Despite the higher complexity and perceived stability of the larger HIF-2 DoHD complexes, we were still able to see disruption of these complexes from all tested small molecule inhibitors, which is especially interesting in the case of KG-279 and KG-548, which had not yet been reported as capable of disrupting HIF-2 dimerization, even in single heterodimers (1).

Finally, our work here sheds some much-needed light on the nature of interactions between CCCs and the HIF-α/ARNT heterodimers. While past work established the importance of these interactions in HIF-driven transcriptional activation via ARNT (10, 40, 55–59), prior work from our lab and others led to different, and conflicting, views as to the structural basis of activator/coactivator recruitment (2, 9, 15, 39). Our complexes here of HIF-1 and HIF-2 bound to different TACC3 fragments resolve this dilemma, showing TACC3 alongside the length of either HIF complex, with what we believe to be the C-terminus of TACC3 directly interacting with the ARNT PAS-B domain based on prior work from our lab (2).. Additional interactions are indicated by cryo-EM (and HDX-MS for HIF-2), showing the involvement of loops on the HIF-α subunit. Notably, the HDX-MS data shows a clear stabilization of the ARNT PAS-A/PAS-B linker upon TACC3 binding; a similar stabilization has been observed with a HIF-2 binding small molecule activator (24), raising the question about similar activation modes despite the differences in activator binding to the HIF heterodimer. The recruitment of TACC3 to a single HIF heterodimer bolsters the idea that HIFs may form markedly different complexes depending on the functional context; in a case where HIFs cannot form dimer-of-heterodimer complexes, we believe that other components, such as CCCs, are recruited to augment HIF function.

We close by underscoring that structural biology – like all experimental methods – has its fundamental strengths and weaknesses. Within HIF biology, the strengths are clear as evidenced by mechanistic understanding of how natural O_2_ regulation is achieved, how disease-associated disruptions have their effects, and how small molecules can be used therapeutically to target those diseases. However, we equally see reminders that single structures of complexes, while extremely useful on their own, may not entirely represent protein assemblies in their true functional contexts, due to experimental liabilities (*e.g.* presence or absence of protein dynamics) or experimental choice (*e.g.* selection of proteins, DNAs, small molecules). Our work here underscores the importance of the latter; with the rich variety of HIF interacting partners and promoters known and yet to be discovered, many discoveries remain ahead.

## Materials and Methods

### Cloning and Expression of HIF Complexes

6xHis-tagged wildtype human HIF-2α (residues 5-361) or HIF-1α (residues 4-361) and untagged human ARNT (residues 91-470) were cloned into a pETDuet-1 vector. HIF-α (6xHis-tagged) and ARNT were coexpressed in BL21-CodonPlus (DE3) *E. coli* (Agilent) in Lysogeny Broth with 0.1 mg/mL ampicillin and 0.034 mg/mL chloramphenicol. Cells were grown at 37°C and induced with 0.1 mM isopropyl-β-D-thiogalactopyranoside (IPTG) upon reaching an OD_600_ of 0.8-1.0, then brought to 16°C to induce overnight (16-18 hr). Cells were harvested by spinning at 4°C and 4658xg for 45 min, resuspended in lysis buffer (50 mM HEPES (pH 7.4), 300 mM NaCl, 3 mM β-mercaptoethanol (BME), 1 mM phenylmethylsulfonyl fluoride (PMSF), 0.1 mg/mL lysozyme and 0.01 U/mL benzonase nuclease), flash frozen and stored at -80°C.

### Cloning and Expression of Coiled-Coil Coactivators

Wildtype human TACC3 (residues 758-838 or residues 788-838 [shTACC3]) or wildtype human CoCoA (residues 407-535) were cloned into a pHis-GB1 vector. CCCs were then expressed in BL21 (DE3) *E. coli* (Agilent) in Lysogeny Broth with 0.1 mg/mL ampicillin. Cells were grown at 37°C and induced with 1 mM isopropyl-β-D-thiogalactopyranoside (IPTG) upon reaching an OD_600_ of 0.8-1.0, then brought to 17°C to induce overnight (16-18 hr). Cells were harvested by spinning at 4°C and 4658xg for 45 min, resuspended in lysis buffer (50 mM Tris (pH 7.4), 150 mM NaCl, 1 mM PMSF 0.1 mg/mL lysozyme and 0.01 U/mL benzonase nuclease), flash frozen and stored at -80°C.

### Cloning and Expression of ARNT PAS-B

Wildtype human ARNT PAS-B (residues 356-470) was cloned into a pHis-GB1 expression vector. ARNT PAS-Bs were then expressed in BL21 (DE3) *E. coli* cells (Agilent) in Lysogeny Broth with 0.1 mg/mL ampicillin. Cells were grown at 37°C and induced with 1 mM isopropyl- β-D-thiogalactopyranoside (IPTG) upon reaching an OD_600_ of 0.8-1.0, then brought to 17°C to induce overnight (16-18 hr). Cells were harvested by spinning down at 4°C and 4658xg for 45 minutes, resuspended in lysis buffer (50 mM Tris (pH 7.4), 150mM NaCl, 2mM BME, 1mM PMSF), flash frozen and stored at -80°C.

### Purification of HIFα-ARNT Heterodimers

Frozen HIF-1 or HIF-2 cell pellets were thawed, lysed by sonication and centrifuged at 47850xg for 45 min. HIF-α-ARNT was purified using an AKTA Püre (Cytiva) by flowing lysate over a 5 mL HisTrap HP (Cytiva), followed by wash buffer (50 mM HEPES (pH 7.4), 300 mM NaCl, 20 mM imidazole, 2 mM BME) with stepwise increasing concentrations of imidazole (20-260 mM) to remove nonspecifically bound protein. 6xHis-tagged HIF-α (bound to ARNT) was eluted with elution buffer (50 mM HEPES (pH 7.4), 300 mM NaCl, 500 mM imidazole, 2 mM BME), mixed with about a 20x molar excess of annealed DNA (20 bp, 51 bp BS1 or 52 bp BS2, IDT) and 5 mM MgCl_2_ and allowed to bind for at least one hour at 4°C. Protein was then concentrated about 10x by dialyzing (3.5 kDa MWCO dialysis tubing, ThermoFisher) against dry polyethylene glycol 35000. HRE-bound HIF-α-ARNT complexes were then further purified through a Superdex 200 increase 10/300 analytical size exclusion column (Cytiva) with the final analytical buffer (50 mM HEPES (pH 7.4), 250 mM NaCl, 5 mM MgCl_2_). Protein samples were used immediately or stored at 4°C for a maximum of 48 hr. 20 bp HRE: GGCTGCGTACGTGCGGGTCGT 51 bp BS1 HRE: AGGGGTGGAGGGGGCTGGGCCCTACGTGCTGTCTCACACAGCCTGTCTGAC 52 bp BS2 HRE: TGGGCCCTACGTGCTGTCTCACACAGCCTGTCTGACCTCTCGACCCTACCGG

### Purification of Coiled-Coil Coactivators

Frozen cell pellets were thawed, lysed by sonication and centrifuged at 47850xg for 45 min. TACC3 or CoCoA were purified using an AKTA Püre (Cytiva) by flowing lysate over a 5 mL HisTrap HP (Cytiva), followed by wash buffer (50 mM Tris (pH 7.4), 150 mM NaCl, 20 mM imidazole) with stepwise increasing concentrations of imidazole (39-106 mM) to remove nonspecifically bound protein. 6xHis-tagged-GB1-CCC (TACC3 or CoCoA) was eluted with elution buffer (50 mM Tris (pH 7.4), 150 mM NaCl, 500 mM imidazole) and diluted 1:10 in TEV buffer (50 mM Tris (pH 7.4), 0.1 mM EDTA) with a 3:1000 addition of TEV protease to cleave the His and GB1-tags from the protein. Cleaved tags were then removed by flowing the protein over a 5 mL HisTrap HP to separate them from the protein of interest. CCC-containing fractions were then concentrated about 15x using a 10 kDa MWCO Millipore centrifugal filter unit (Sigma-Aldrich) at 1200xg and 4°C. Concentrated protein was further purified through a Superdex 200 increase 16/40 HiScale size exclusion column (Cytiva) with the final analytical buffer (50 mM HEPES (pH 7.4), 50 mM NaCl, 5 mM MgCl_2_). Protein samples were used immediately or stored at 4°C for a maximum of 7 days.

### Purification of ARNT PAS-B

Frozen cell pellets were thawed, lysed by sonication and centrifuged at 47850xg for 45 min. ARNT PAS-B was purified using an AKTA Püre (Cytiva) by flowing lysate over a 5 mL HisTrap HP (Cytiva), followed by wash buffer (50 mM Tris (pH 7.4), 150 mM NaCl, 20 mM imidazole, 1 mM DTT) with stepwise increasing concentrations of imidazole (39-106 mM) to remove nonspecifically bound protein. 6xHis-tagged-GB1-ARNT PAS-B was eluted with elution buffer (50 mM Tris (pH 7.4), 150 mM NaCl, 500 mM imidazole, 1 mM DTT) and diluted 1:10 in TEV buffer (50 mM Tris (pH 7.4), 0.1 mM EDTA) with a 3:1000 addition of TEV protease to cleave the His and GB1-tags from the protein. Cleaved tags were then removed by flowing the protein over a 5 mL HisTrap HP to separate them from the protein of interest. ARNT PAS-B-containing fractions were then concentrated about 15x using a 10 kDa MWCO Millipore centrifugal filter unit (Sigma-Aldrich) at 1200xg and 4°C. Concentrated protein was further purified through a Superdex 75 increase 16/40 HiScale size exclusion column (Cytiva) with the final analytical buffer (50 mM HEPES (pH 7.4), 15 mM NaCl, 1 mM DTT). Protein samples were used immediately or stored at -80°C.

### Cryogenic Electron Microscopy

Purified HIF complexes were concentrated to a final concentration of 10-20 µM (1-2 mg/mL) using 10 kDa Pierce Concentrators (ThermoFisher) at 4°C and 1000xg. TACC3 (residues 788- 838) was mixed with the HIF-2 complexes to a final concentration of 70-150 μM and allowed to bind for ∼20 min on ice. 3.5 µL of sample was applied to Quantifoil Au 300 R1.2/1.3 with Ultrathin Carbon EM grids (Electron Microscopy Sciences) and plunge frozen using a Vitrobot 3 (ThermoFisher) at 100% humidity and 4°C. Frozen grids were then analyzed at the New York Structural Biology Center (NYSBC) Members Electron Microscopy Center (MEMC) using a TFS Titan Krios transmission electron microscope (ThermoFisher). Micrographs were collected at a pixel size of 0.844 Å/px at an exposure rate of 58.94 e^-^/Å^2^/frame with a defocus range of 0.8-2.0 μm. Particles were picked from micrographs and extracted to a box size of 400x400 pixels using Warp (41) and analyzed using CryoSPARC (42). Particles were typically sorted through iterative rounds of 2D classification, *ab initio* reconstruction, and heterogeneous, homogenous and non-uniform refinement.

### Amplified Luminescent Proximity Homogenous Assay (Alpha) Screen

HIF-2 complexes with C-terminally FLAG-tagged (DYKDDDDK) ARNT subunits were purified and bound to either 20 bp HRE or 51 bp HRE/HAS biotinylated DNA fragments (IDT). 300 nM of the resulting complexes were incubated with 20 μg/mL streptavidin Alpha donor beads (Revvity), 20 μg/mL anti-FLAG Alpha acceptor beads (Revvity), and varying concentrations of KG-548, KG-279, compound 2 and PT2385 with a final DMSO concentration of 1% (v/v) and a final tween-20 concentration of 0.05% (v/v). Automation of sample preparation was accomplished using an Opentrons OT-2 liquid handler (Opentrons Labworks Inc.). Samples were allowed to incubate one hour at 25°C in the dark before being analyzed on a SpectraMax i3 plate reader (Molecular Devices) equipped with an AlphaScreen 384 STD cartridge (Molecular Devices). Data was collected using SoftMax Pro (Molecular Devices) and plotted with Grace. All samples were run in triplicate.

### Crystallization and Crystallographic Structure Determination of TACC3(758–838)

12.5 mg/mL of purified human TACC3(758–838) and 9.3 mg/mL of human ARNT PAS-B(356–470) were mixed in 50 mM HEPES, pH 7.4, 50 mM NaCl, 50 mM MgCl_2_, and 5% glycerol crystallized in 1 M imidazole, pH 7 and 50% (v/v) MPD via the sitting-drop vapor diffusion method. The condition was found via high-throughput screening using NeXtal Protein Complex sparse matrix crystallization screen (NeXtal Technologies; condition H9, the drop contained equal volumes of protein solution and mother liquor). Crystals were harvested directly from the screen and cryoprotected with LV CryoOil (MiTeGen) and flash-cooled in liquid N_2_ prior to data collection. Data were collected at the National Synchrotron Light Source II on the AMX (17-ID- 1) beamline at Brookhaven National Laboratory. Data were processed using the autoPROC tool box (60) resulting in a 2.28 Å data set. While electron density for ARNT PAS-B was not observed, sufficient density for TACC3 residues 780-836 was visible to allow the structure to be determined by molecular replacement with Phaser (61) using PDB entry 4PKY (chains B and C) as the starting search model. Several cycles of refinement were conducted using Coot (62) and Phenix.refine (63). Data collection, processing and refinement statistics are provided in **Table S2**. Coordinates for TACC3(758–838) have been deposited in the RCSB with accession number 9YDY, including residues 780-836 on one chain and 782-836 on the other.

### Microscale Thermophoresis

Purified HIF complexes bound to carboxyfluorescein (FAM) labeled HRE (IDT) were diluted to a final concentration of 1μM fluorophore, with an addition of 0.05% tween-20. ARNT PAS-B proteins (20 μM) were instead labeled with fluorophore at free amino sites using an ALEXA Fluor 488 N-hydroxysuccinimide (NHS)-ester (Thermo Fisher) by first exchanging into pH 8.2 analytical buffer, allowing incubation with 50 μM of ALEXA 488 NHS ester for 1 hour at 4°C in the dark, then exchanging back into analytical buffer to remove excess dye. Final protein concentration is calculated using the following equation (where A_280_ is the absorbance reading at 280 nm, A_495_ is the absorbance reading at 495 nm, and ε_280_ is the extinction coefficient of the protein at 280 nm) and adjusted to 1 μM.

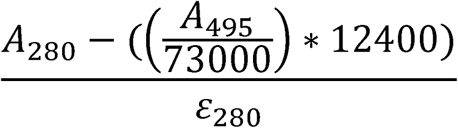

A serial dilution of shTACC3 was generated in TACC3 buffer (50 mM HEPES (pH 7.4), 50 mM NaCl, 5 mM MgCl_2_). MST samples were generated as a 1:1 (v/v) mix of HIF complex (fixed 500 nM final concentration) to each concentration of the TACC3 serial dilution. Samples were drawn into standard MST capillaries (NanoTemper) and loaded into a Monolith NT.115 (NanoTemper) for binding analyses. Data were obtained using MO.Control (NanoTemper) and analyzed using PALMIST and GUSSI (64).

### Structural modeling of HIF complexes

AlphaFold-generated structures were flexibly fit into refined cryo-EM density maps using ChimeraX (65) and ISOLDE (66) to apply distance restraints. For the HIF-2 dimer of heterodimers, we flexibly fit our cryo-EM structure of human HIF-2 into the obtained cryo-EM density. Structures were then further refined using PHENIX (63) to perform global real space refinement and Coot (62) to perform atomic refinement and remove unresolved regions. Structural validation was performed using MolProbity and comprehensive validation in PHENIX (63). For HIF-1 and HIF-2/TACC3 complexes, shTACC3 fragments were fit into corresponding cryo-EM maps (with the HIF complex docked and fixed) using the Dock in Map function of PHENIX (63) and assuming that the TACC3 C-terminus is closest to ARNT PAS-B (10). We emphasize that the HDX-MS data collected on HIF-2/TACC3 complex was not used to guide this model generation, as it was saved for post-modeling validation. Coordinates for HIF-1 and HIF-2 complexes were deposited with RCSB under the following accession numbers: 9OF0 (HIF-2 heterodimer/HRE; EMDB EMD-70416), 9OF2 (HIF-2 dimer of heterodimers/BS1 HRE/HAS; EMDB EMD-70418), 9OFU (HIF-1 dimer of heterodimers/BS2 HRE/HAS; EMDB EMD-70443). Coordinates for models of TACC3/HIF-1/HRE and TACC3/HIF-2/HRE complexes are deposited at PDB-IHM under the accession numbers 9A9Z (TACC3/HIF-1) and 9A9Y (TACC3/HIF-2).

### Size Exclusion Chromatography Coupled with Multi-Angle Light Scattering (SEC-MALS)

Purified HIF complexes were concentrated to a final concentration of 15-20 µM (1-2 mg/mL) using 10 kDa Pierce Concentrators (ThermoFisher) at 4°C and 1000xg. Samples were then filtered through an Ultrafree 0.1 µm centrifugal filter (Millipore) at 4°C and 1000xg. 500 µL of sample were passed through a Superdex 200 10/300 size exclusion column (Cytiva) with the HIF analytical buffer (50 mM HEPES (pH 7.4), 250 mM NaCl, 5 mM MgCl_2_) before passing through a DAWN HELEOS II MALS detector and an Optilab T-rEX Refractive Index Detector (Wyatt/Waters) using a P-920/UPC-900 AKTA FPLC System (Cytiva). SEC-MALS data was obtained and analyzed using ASTRA (Wyatt/Waters) and plotted in R.

### Size Exclusion Chromatography for HIF-1/HIF-2 Complex Determination

Human HIF-1α (residues 4-361) and ARNT (residues 91-470) were cloned into a pHis-MBP vector (67) and expressed as detailed above for HIF-complexes. MBP-tagged HIF-1 and HIF-2 were separately lysed through sonication and centrifuged for 45 min at 47850xg and 4°C. Lysates were then combined and purified together using an AKTA Püre (Cytiva) by flowing lysate over a 5 mL HisTrap HP (Cytiva), followed by wash buffer (50 mM HEPES, 300 mM NaCl, 20 mM imidazole, 2 mM BME) with stepwise increasing concentrations of imidazole (20 mM, 68 mM, 164 mM, 260 mM) to remove nonspecifically bound protein. Mixed proteins were then eluted with elution buffer (50 mM HEPES (pH 7.4), 300 mM NaCl, 500 mM imidazole, 2 mM BME), then mixed with about a 20x molar excess of 51 bp BS1 HRE, 51 bp BS1 DNA with a mutated HRE-box, or 51 bp BS1 DNA with a mutated HAS-box (IDT) and 5 mM MgCl_2_ and allowed to bind at 4°C overnight (10-16 hr). Protein was then concentrated about 10x by dialyzing (3.5 kDa MWCO dialysis tubing, ThermoFisher) against dry polyethylene glycol 35000. DNA-bound MBP-HIF-1/HIF-2 complexes were further purified through a Superdex 200 increase 16/40 HiScale (Cytiva) with the final analytical buffer (50 mM HEPES, (pH 7.4) 250 mM NaCl, 5 mM MgCl_2_) to separate complexes via size exclusion chromatography. Complex components were then visualized using gel electrophoresis. SEC profiles were plotted in R.

Mutant HRE-box BS1 DNA: AGGGGTGGAGGGGGCTGttaaaTgatatCTGTCTCACACAGCCTGTCTGAC

Mutant HAS-box BS1 DNA: AGGGGTGGAGGGGGCTGGGCCCTACGTGCTGTCgatatgtcgaCtTCTGAC

### Hydrogen Deuterium Exchange Coupled with Mass Spectrometry (HDX-MS)

Purified 20 bp HRE-bound HIF-2 was incubated with shTACC3 for one hour in about a 3:1 molar ratio of TACC3 to HIF-2 (74 μM shTACC3 and 24 μM HIF-2) or an equivalent volume of TACC3 buffer (50 mM HEPES, 50 mM NaCl, 5 mM MgCl_2_). Using a LEAP HDX Automation Platform (Trajan Automation), the protein mixture was deuterated in D_2_O buffer (50 mM HEPES, 50 mM NaCl, 5 mM MgCl_2_, solvated in 100% D_2_O) for varying timepoints (0 s, 50 s, 300 s, 1000 s, 3000 s, 5000 s) after which the deuterated protein mixture was mixed with quench buffer (3 M guanidinium HCl, 3% acetonitrile, 1% formic acid) at 0°C. Samples were then loaded onto a Waters Enzymate BEH Pepsin Column for proteolysis followed by a C18 analytical column (Hypersil Gold, 50 mm length × 1 mm diameter, 1.9 μm particle size, ThermoFisher Scientific) before being loaded onto a maXis-II ETD ESI-qTOF mass spectrometer (Bruker) for fragment mass analysis. Timepoints both with and without TACC3 present were completed in triplicate. Raw HDX data was analyzed using Bruker Compass Data Analysis 5.3 and Biotools 3.2 software and further analyzed with HDexaminer (Sierra Analytics/Trajan Automation) to calculate relative exchange rates before being plotted in R. Samples were independently run in triplicate.

### Reporter assays

Luciferase reporter assays were performed with transiently transfected HEK293T cells. Cells were plated onto 48-well plates at 100,000 cells/well. Twenty-four hours later, cells were transfected in triplicate with Lipofectamine 2000 (Invitrogen). Each well received 33 ng reporter plasmid under control of wild-type or mutant human 3’ erythropoietin (Epo) enhancer fused to a firefly luciferase cassette (pGL3-Basic, Promega), 200 ng empty expression plasmid (pIRES- hrGFP-2a, Stratagene), and 100 ng plasmid expressing oxygen insensitive “PPN” HIF-1α or HIF-2α variants, which contain mutations in two proline and one asparagine residue that are normally subject to O_2_-dependent post-translational modifications (68). At 24 hr post- transfection, cells were rinsed one time with PBS (ThermoFisher), followed by lysis with 50 μL core luciferase buffer (30 mM tricine (pH 7.8), 8 mM magnesium acetate, 0.2 mM EDTA, 1% Triton X-100, and 100 mM 2-mercaptoethanol). Luciferase assays were performed by adding 50 μL substrate buffer (core buffer supplemented with 2.5 mM MgCl_2_, 1.5 mM ATP, 0.5 mM coenzyme A, and 0.5 mM D-luciferin) to 10 μL cell lysate and immediately reading luminescence on a CLARIOstar Plus plate reader (BMG Labtech). Long oligos with Nhe I and Xho I restriction sites at the 5’ and 3’ ends, respectively, were used to position the 3’ human Epo enhancer upstream of a minimal mouse Epo promoter. Sequences of the 3’ human Epo enhancer sequences used in this study are indicated below (mutations and restriction sites in lowercase):

WT: gctagcAGATCTGGGAAACGAGGGGTGGAGGGGGCTGGGCCCTACGTGCTGTCTCACACAGCCTGTCTGACCTCTCGACCCTACCGGGCCTGAGGCCACAAGCTCTGCCTACGCT GGTCAATAAGGTGGCTCCATTCAAGGCCTCACCGCAGTAAGGCAGCTGCCAACCCTcgag

GC-rich mutant: gctagcAGATCTGGGAAACGAGGctaGaAttcctCTGGGCCCTACGTGCTGTCTCACACAGCCTGTCTGACCTCTCGACCCTACCGGGCCTGAGGCCACAAGCTCTGCCTACGCTGGTCAATAAGGTGGCTCCATTCAAGGCCTCACCGCAGTAAGGCAGCTGCCAACCCTcgag

HRE mutant: gctagcAGATCTGGGAAACGAGGGGTGGAGGGGGCTGttaaaTgatatCTGTCTCACACAGCCTGTCTGACCTCTCGACCCTACCGGGCCTGAGGCCACAAGCTCTGCCTACGCTGGTCAATAAGGTGGCTCCATTCAAGGCCTCACCGCAGTAAGGCAGCTGCCAACCCTcgag

HAS mutant: gctagcAGATCTGGGAAACGAGGGGTGGAGGGGGCTGGGCCCTACGTGCTGTCgatatgtcgaCtTCTGACCTCTCGACCCTACCGGGCCTGAGGCCACAAGCTCTGCCTACGCTGGTCAATAAGGTGGCTCCATTCAAGGCCTCACCGCAGTAAGGCAGCTGCCAACCCTcgag

## Acknowledgements

We thank Dan Lee and Julia Simon for technical assistance, Soumendu Boral and other members of the Gardner laboratory as well as Rick Bruick, Uttam Tambar, Qing Zhang, and Derek Tan for helpful discussions. This work was supported by NIH grants U54 CA132378 and U54 CA137788 (to K.H.G.), Mathers Foundation grant MF-2112-02288 (to K.H.G. and J.A.G.), and funds from the Department of Veterans Affairs I01BX000446 (to J.A.G.). Cryo-EM data collection was performed at the Members Electron Microscopy Center at the New York Structural Biology Center, with major support from the Simons Foundation (SF349247). This research used the AMX and FMX beamlines of the National Synchrotron Light Source II, a U.S. Department of Energy (DOE) Office of Science User Facility operated for the DOE Office of Science by Brookhaven National Laboratory under Contract No. DE-SC0012704.

