## Supporting Information for "Context-Dependent Variability Of HIF Heterodimers Influences Interactions With Macromolecular And Small Molecule Partners"

Joseph D. Closson[1,2], Xingjian Xu[1,2], Meiling Zhang[1], Tarsisius T. Tiyani[1,2], Leandro Pimentel Marcelino[1,3], Eta A. Isiorho[1], Jason S. Nagati[4,5], Joseph A. Garcia[4,5], Kevin H. Gardner[1,3,6]

[1]: Structural Biology Initiative, CUNY Advanced Science Research Center, New York, NY 10031

[2]: Ph.D. Program in Biochemistry, The Graduate Center – City University of New York, New York, NY 10016

[3]: Department of Chemistry and Biochemistry, City College of New York, New York, NY 10031

[4]: Department of Medicine, Columbia University Irving Medical Center, New York, NY 10032

[5]: Departments of Medicine and Research & Development, James J. Peters Veterans Affairs Medical Center, Bronx, NY 10468

[6]: Ph.D. Programs in Biochemistry, Biology, and Chemistry, The Graduate Center – City University of New York, New York, NY 10016

Contents:


- Supporting Figures S1-S11
- Supporting Tables S1-S4
- Supporting References

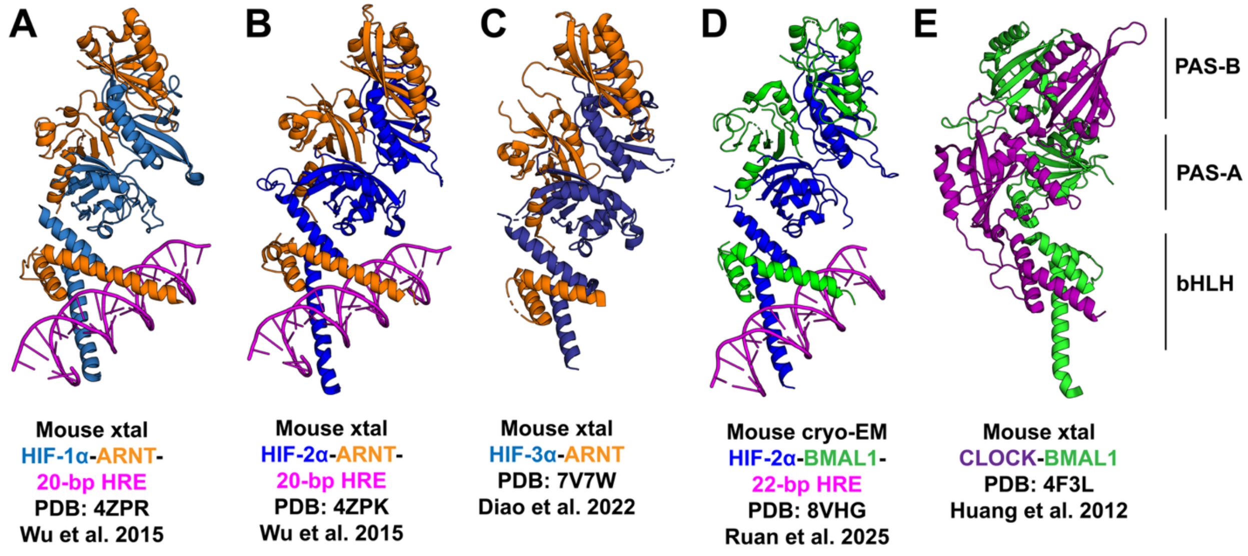


**Figure S1. Existing Structures of HIFs and HIF Homologs.** Crystal structures of murine HIF-1 (A) (1), HIF-2 (B) (1) and HIF-3 (C) (2) show highly similar domain architectures between HIF isoforms. A cryo-EM structure of HIF-2α complexed with BMAL1 (D) (3) shows a similar domain arrangement to the HIF isoforms than CLOCK-BMAL1, implying the HIF-α subunit is a major contributor to domain arrangement. A crystal structure of murine CLOCK-BMAL1 (E) (4) shows similar bHLH and PAS tertiary structures, but in a reorganized quaternary arrangement of the PAS domains from analogous HIF complexes.

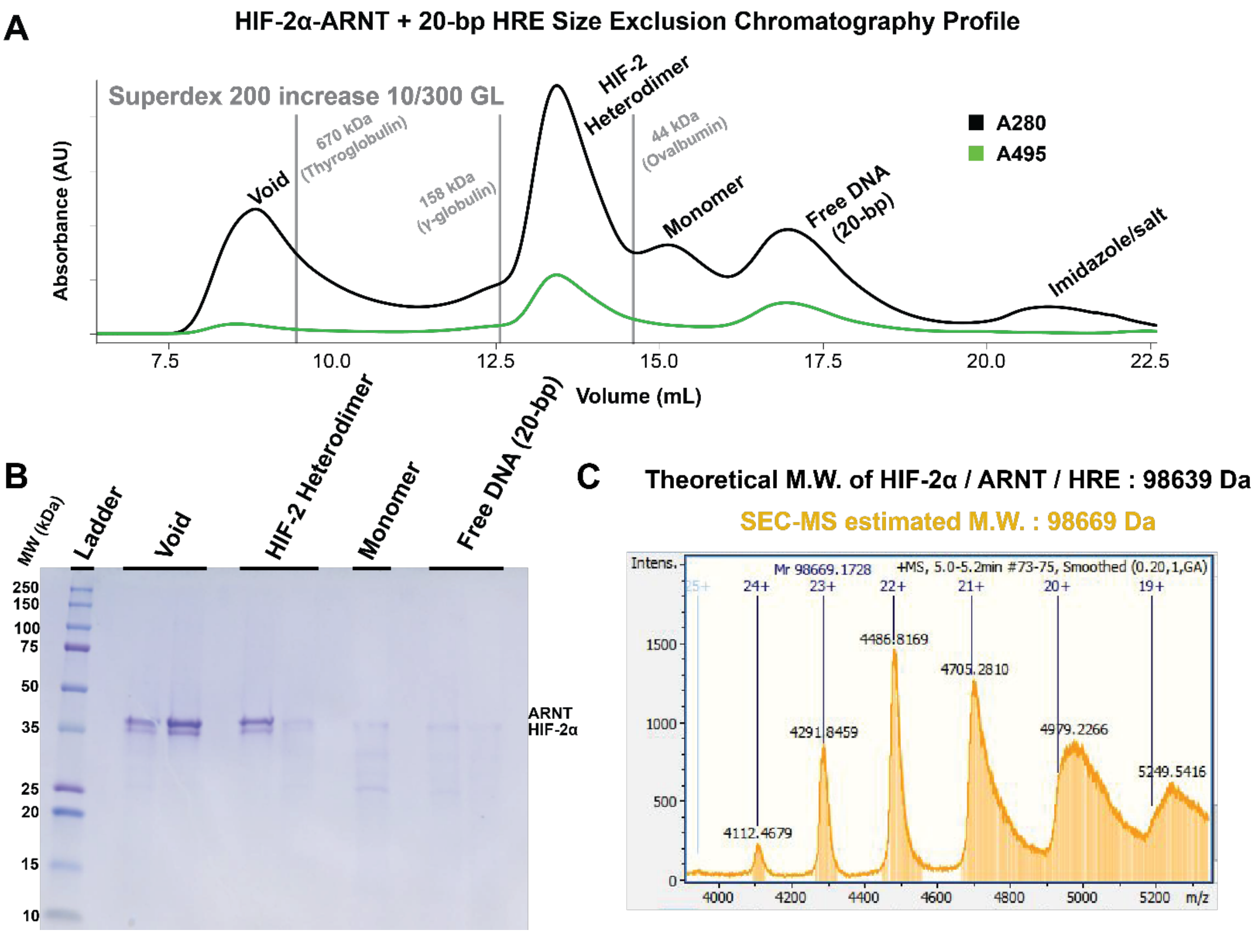


**Figure S2. Characterization of the HIF-2/20-bp DNA Complex.** (A) Superdex 200 increase 10/300 GL purification profile of HIF-2α-ARNT bound to 20-bp FAM-labeled HRE. Absorbance at 280 nm implies presence of protein, while absorbance at 495 nm implies the presence of FAM-labeled HRE, which is seen in both the heterodimer and DNA peaks. (B) SDS-PAGE of respective peaks in size exclusion chromatography (from panel A) shows an abundance of HIF-2α and ARNT (HIF-2α: 42.6 kDa, ARNT: 43.0 kDa) in both void (unfolded) and HIF-2 heterodimer peaks and a small amount of either subunit in the expected monomer peak, along with small molecular weight proteins that are likely remnants from nickel purification. (C) Size exclusion chromatography-mass spectrum of HIF-2 bound to 20-bp DNA. Molecular weight is determined to be 98.669 kDa compared to a theoretical weight of 98.639 kDa.

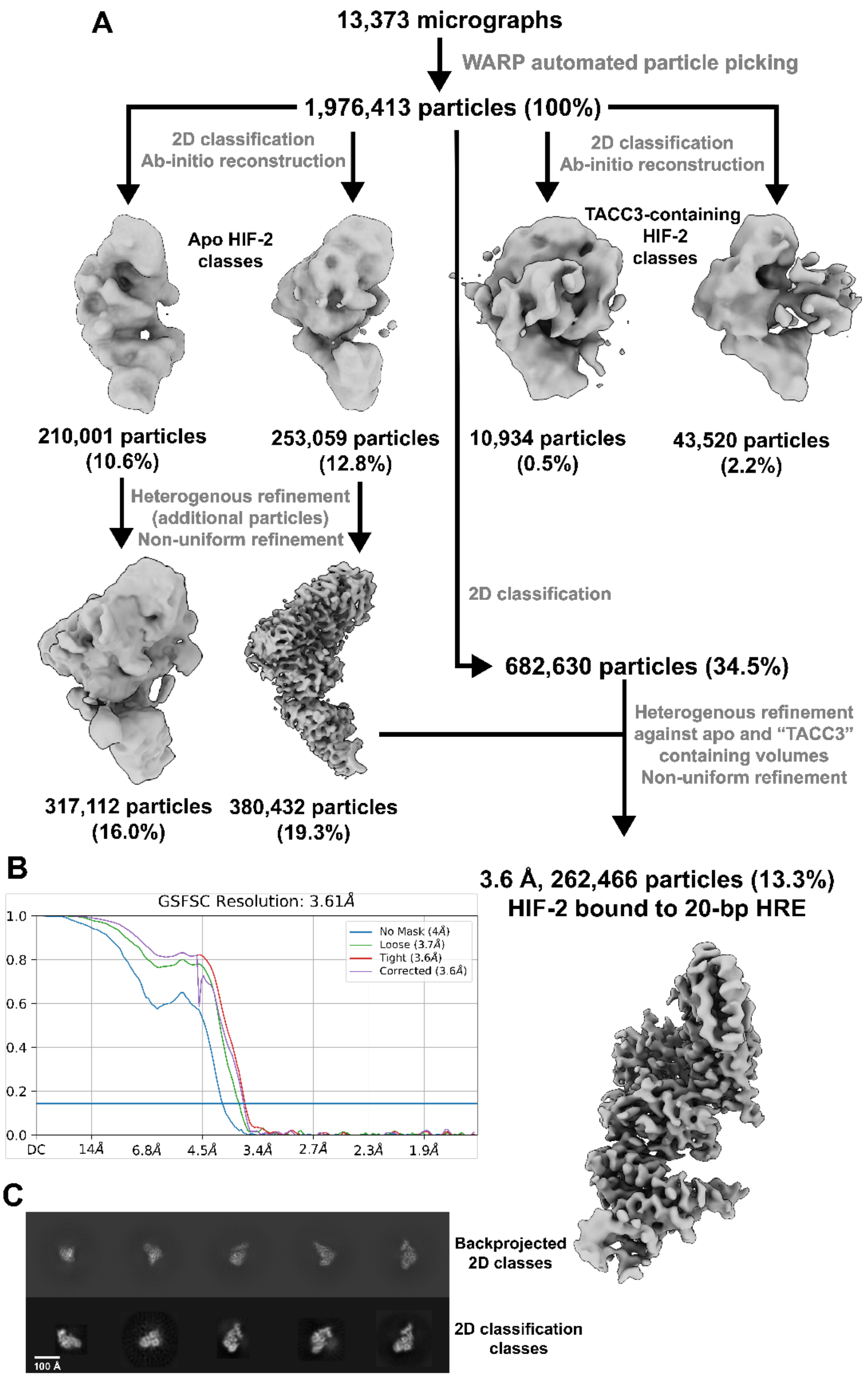


**Figure S3. HIF-2/20-bp DNA Cryo-EM Processing Workflow.** (A) Particles picked from micrographs using WARP (5) were sorted via CryoSPARC (6) into two groups of 3D classes using several rounds of 2D classification and *ab initio* reconstruction, either being apo HIF-2 or containing density corresponding to TACC3. Apo classes were further refined through heterogeneous and non-uniform refinement to a 3.61 Å cryo-EM map containing 262,466 particles, 13.3% of the original particles picked. (B) Fourier shell correlation curve of the final 3.61 Å HIF-2 cryo-EM map obtained from CryoSPARC. (C) Theoretical 2D classes (top) were back-projected from the final HIF-2 cryo-EM map and are comparable to experimental 2D classes obtained from initial particle organization (bottom). Scale bar represents 100 Å.

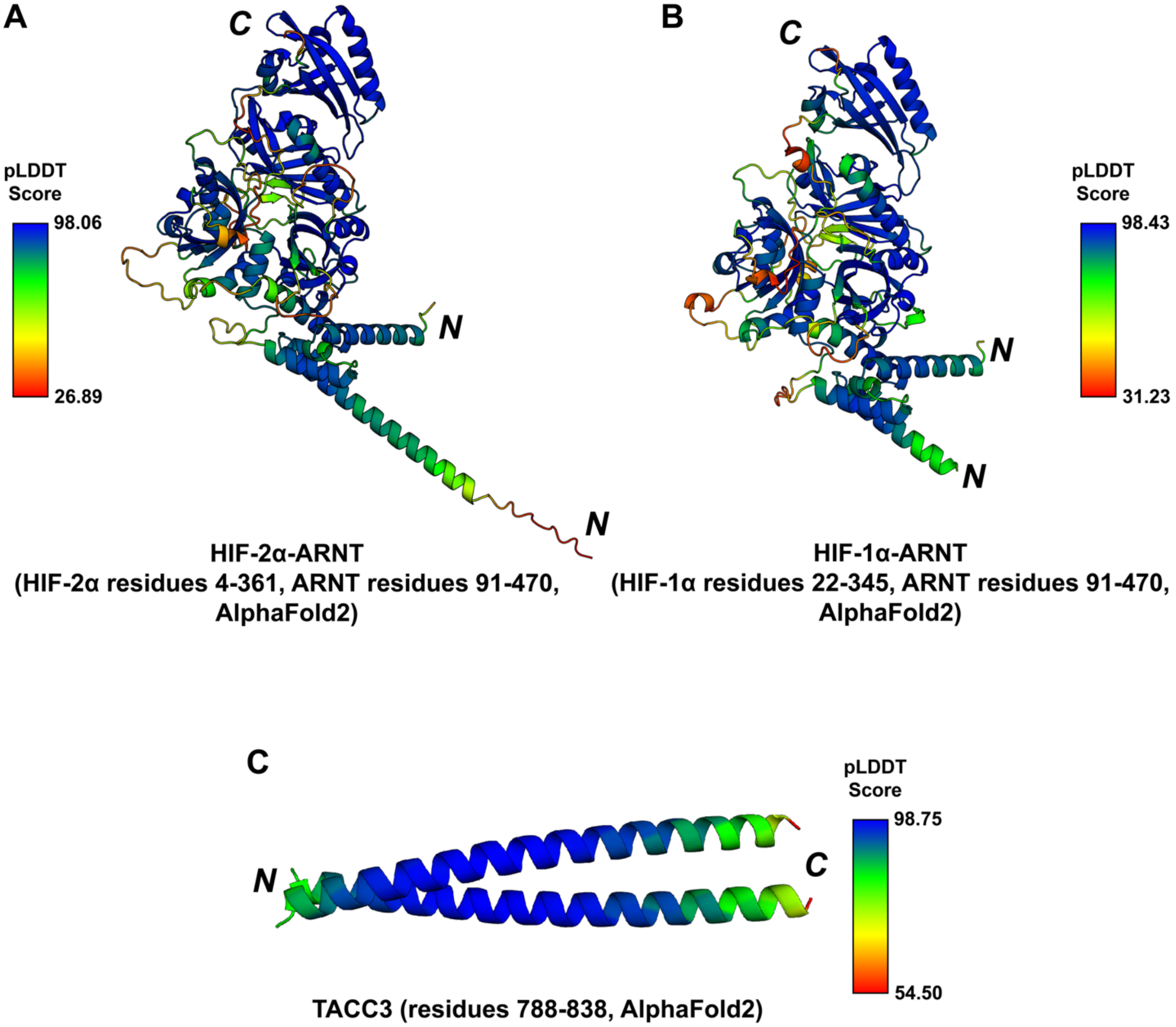


**Figure S4. AlphaFold Models of HIF and TACC3 Proteins Used In Modeling.** AlphaFold 2 (7) generated models of HIF-2 (A), HIF-1 (B) and TACC3 (C) for initial fitting into cryo-EM maps. pLDDT scores were mapped onto the ribbon diagram to assess confidence of the generated model. N and C-termini are indicated for each protein chain.

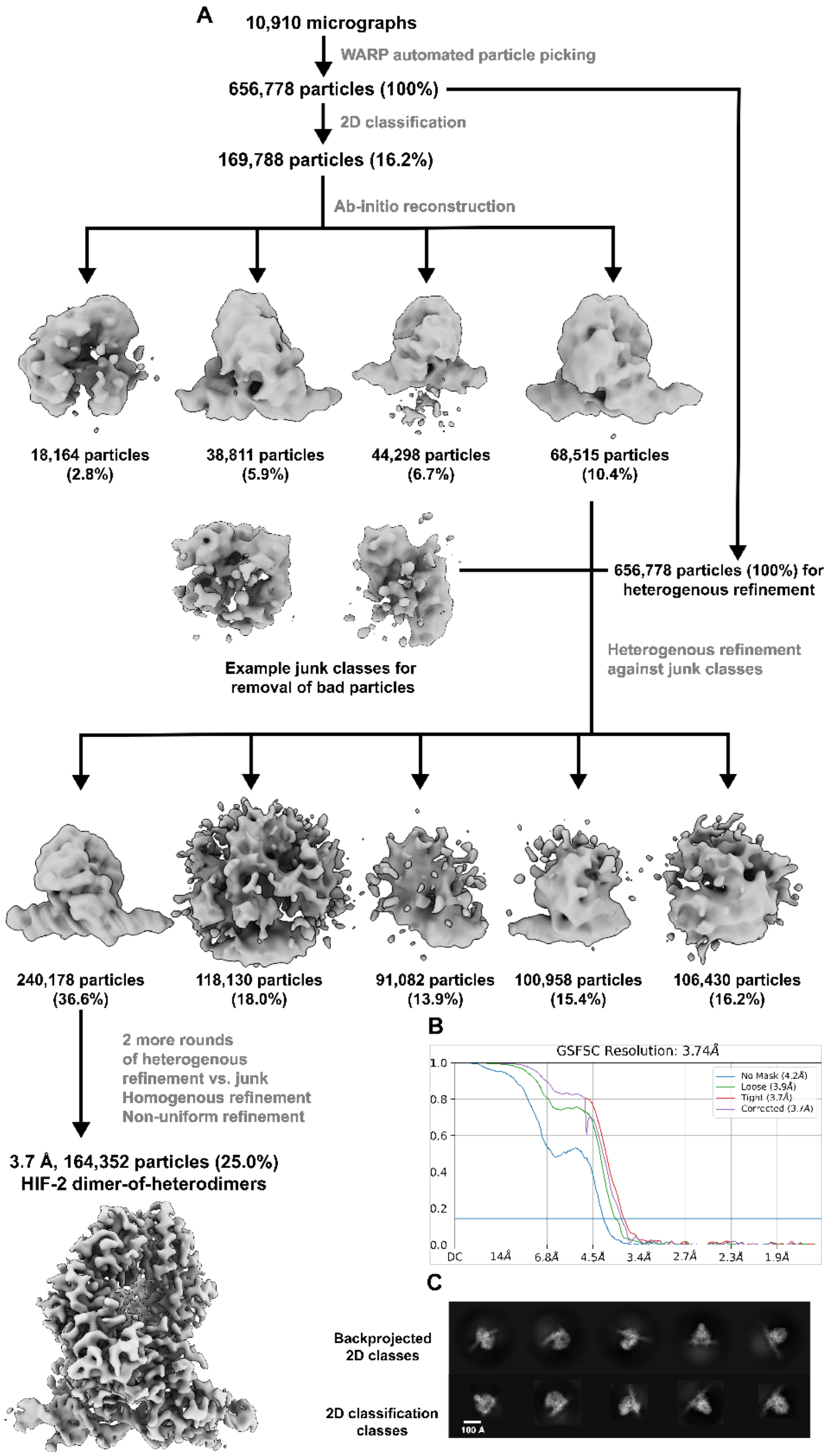


**Figure S5. HIF-2/51-bp BS1 DNA Cryo-EM Processing Workflow.** (A) Particles picked from micrographs using WARP (5) were sorted into several 3D classes using 2D classification and ab-initio reconstruction via CryoSPARC (6). Resulting 3D classes were further refined by sorting particles into "good" and "junk" classes using several rounds of heterogenous refinement and non-uniform refinement, resulting in a 3.74 Å cryo-EM map containing 164,352 particles, 25.0% of the original particles picked. (B) Fourier shell correlation curve of the final 3.74 Å HIF-2/HIF-

2 cryo-EM map obtained from CryoSPARC. (C) Theoretical 2D classes (top) were back-projected from the final DoHD cryo-EM map and are comparable to 2D classes obtained from initial particle organization (bottom). Scale bar represents 100 Å.

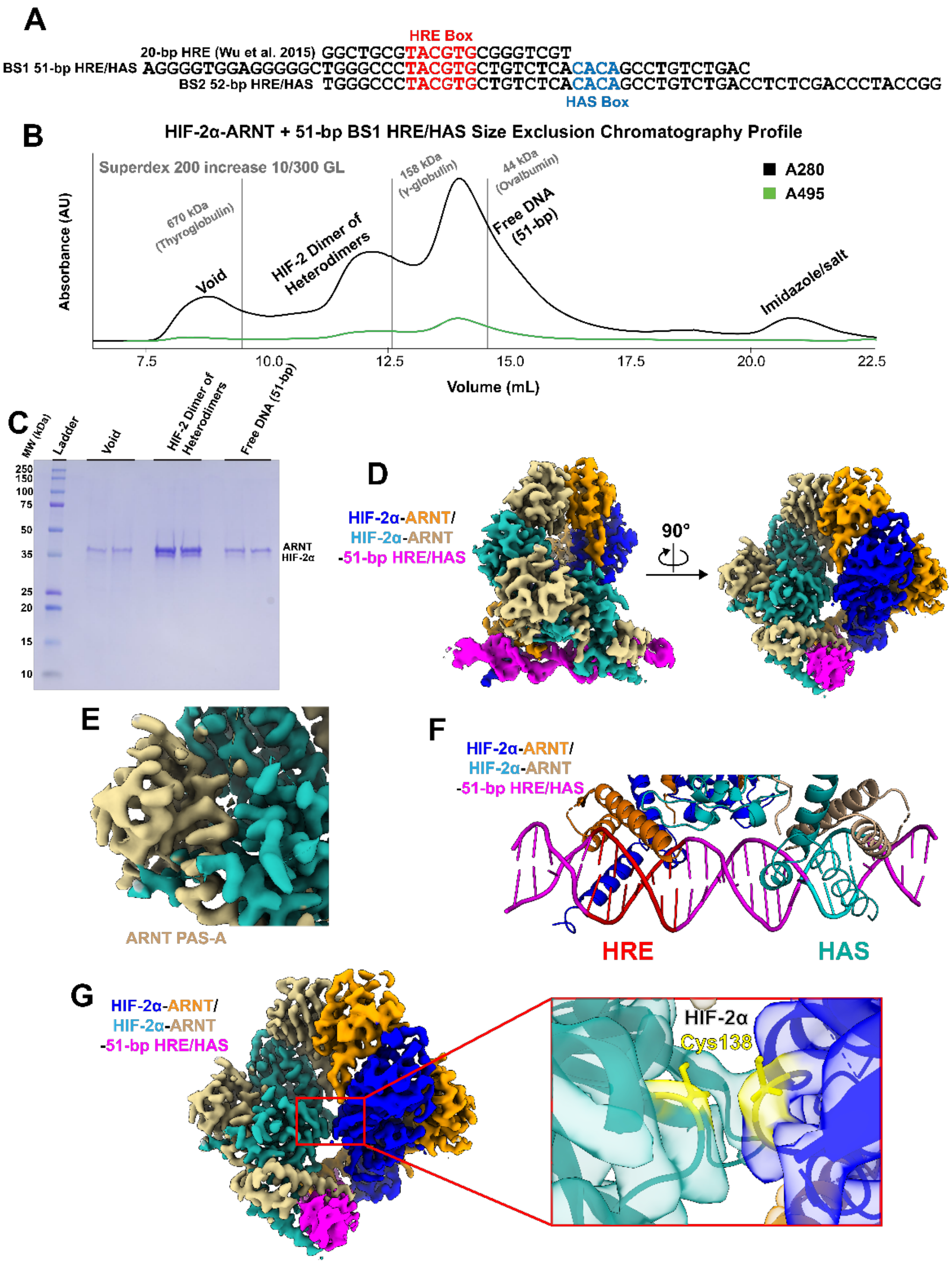


**Figure S6. Characterization of the HIF-2/51-bp BS1 DNA Complex.** (A) DNA sequences used in assembly of HIF-2 complexes. (B) Superdex 200 increase 10/300 GL purification profile of HIF-2α-ARNT bound to 51-bp BS1 FAM-labeled HRE/HAS. Absorbance at 280 nm implies presence of protein, while absorbance at 495 nm implies the presence of FAM-labeled HRE/HAS, which is seen in both the dimer-of-heterodimer and DNA peaks. (C) SDS-PAGE of respective peaks from size exclusion chromatography (from panel B) confirm the presence of both HIF-2α and ARNT (HIF-2α: 42.6 kDa, ARNT: 43.0 kDa) in the void (unfolded) and dimer of heterodimer peaks, with a substantive amount in the free DNA peak. Given the similar elution volume to HIF-2 bound to 20-bp HRE (Fig. S2B), this peak likely also contains single heterodimer bound to HRE, or monomeric HIF-2α or ARNT. (D) Human HIF-2/51-bp BS1 HRE/HAS complex (HIF-2 dimer-of-heterodimers) cryo-EM map with EM density colored according to associated subunits. (E) Cryo-EM map zoomed into the region corresponding to the ARNT PAS-B domain of the secondary (HAS-box bound) heterodimer, this domain is well resolved in HIF-2. (F) DNA-bound bHLH of the HIF-2 dimer-of-heterodimers structure depicting heterodimers separately bound to the HRE and HAS-boxes. (G) Cryo-EM map of the HIF-2 dimer-of-heterodimers zoomed into the HIF-2α PAS-A domains. Cys138 residues of the HIF-2α PAS-A domains appear proximal, but do not show density supporting a disulfide bond.

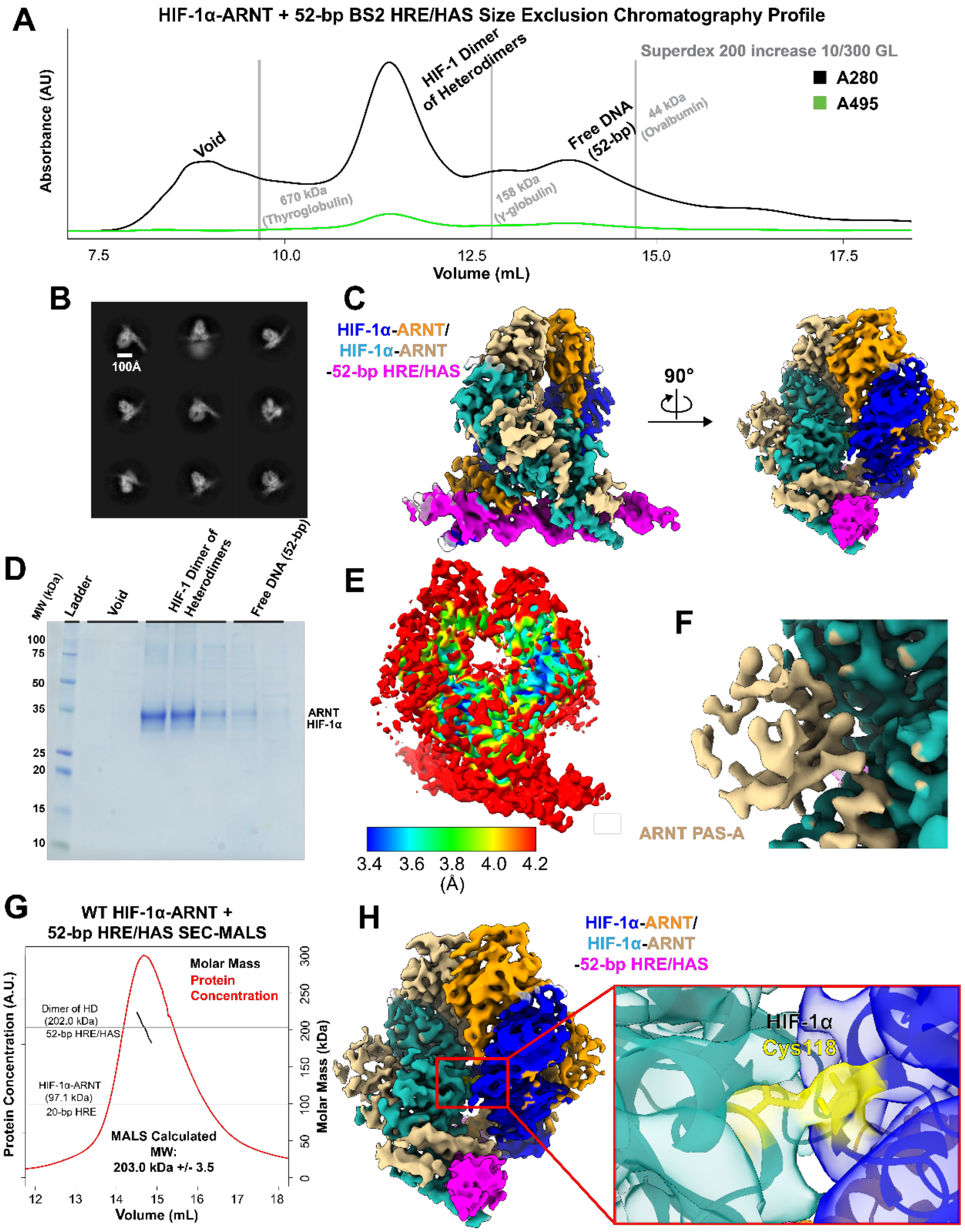


**Figure S7. Characterization of the HIF-1/52-bp BS2 DNA Complex.** (A) Superdex 200 increase 10/300 GL purification profile of HIF-1α-ARNT bound to 52-bp BS2 HRE/HAS. Absorbance at A280 suggests the presence of protein, while absorbance at A495 suggests the presence of FAM-labeled HRE/HAS which is seen in both the dimer-of-heterodimer and free DNA peaks. (B) 2D classes of human HIF-1/52-bp BS2 HRE/HAS complex (HIF-1 dimer-of-heterodimers) cryo-EM particles. Classes are very similar in appearance to HIF-2 dimer-of-heterodimers classes. Scale bar represents 100 Å. (C) Human HIF-1 dimer-of-heterodimers cryo-EM map with EM density colored according to associated subunits. (D) SDS-PAGE of respective peaks from size exclusion chromatography (from panel A) shows the presence of HIF-

1α and ARNT (HIF-1α: 42.9 kDa, ARNT: 43.0 kDa) in the void (unfolded) and dimer of heterodimer peaks, with a small amount in the free DNA peak, which likely correlates to monomeric HIF-1α or ARNT. (E) Local resolution mapped onto the HIF-1 dimer-of-heterodimers cryo-EM map. (F) Cryo-EM map zoomed into the ARNT PAS-B domain of the secondary (HAS-box bound) heterodimer. The resolution of this domain is notably worse than the ARNT PAS-B domains of the HIF-2 dimer-of-heterodimers. (G) SEC-MALS profile of the wildtype HIF-1 dimer-of-heterodimers complex showing the expected molecular weight of the dimer-of-heterodimers complex (203.0±3.5 kDa). (H) Cryo-EM map of the HIF-1 dimer-of-heterodimers zoomed into the HIF-1α PAS-A domains. Cys118 residues of the adjacent HIF-1α PAS-A domains appear proximal and have density spanning them, suggesting the formation of a disulfide bond.

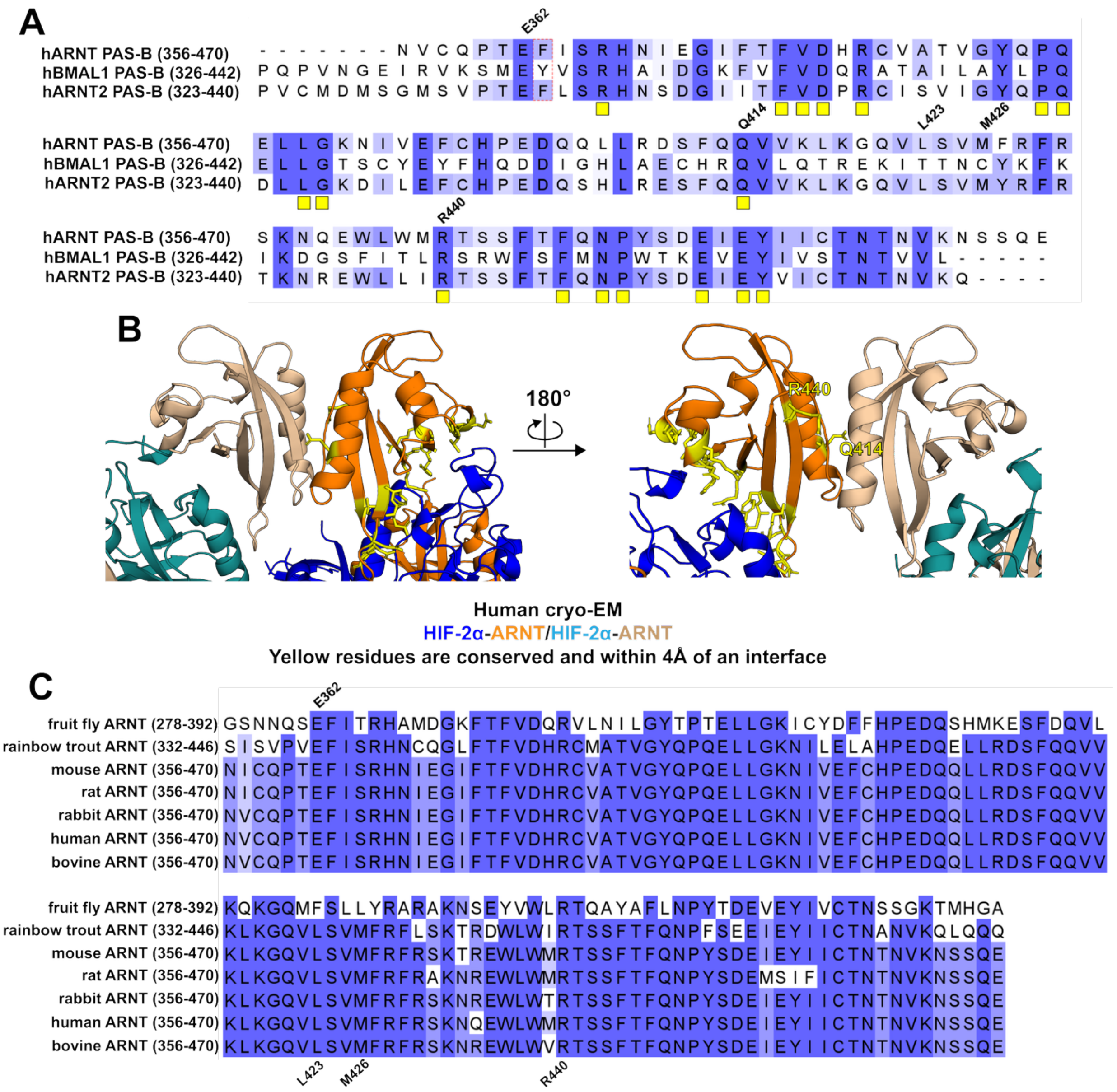


**Figure S8. Assessment of Conserved ARNT Interaction Interfaces.** (A) Sequence conservation of residues between the PAS-B domains of human ARNT homologs human ARNT2 and human brain and muscle ARNT-like 1 (BMAL1). Dark blue residues correspond to residues conserved across the three homologs, while lighter blue residues are partially conserved. Yellow boxes are used to highlight residues located within 4 Å of an interface on the HIF-2 dimer-of-heterodimers. (B) Conserved residues within 4 Å of an interface were mapped (yellow) on the primary ARNT PAS-B domain of the HIF-2 dimer-of-heterodimers. ARNT R440 and Q414 are conserved and located within the interface of the ARNT PAS-B domains on the dimer-of-heterodimer complexes. (C) Sequence alignment of special ARNT homologs (fruit fly [*Drosophila melanogaster*], rainbow trout [*Oncorhynchus mykiss*], mouse [*Mus musculus*], rat [*Rattus norvegicus*], rabbit [*Oryctolagus cuniculus*], human [*Homo sapiens*], bovine [*Bos taurus*]) shows highly conserved ARNT across all species but fruit fly.

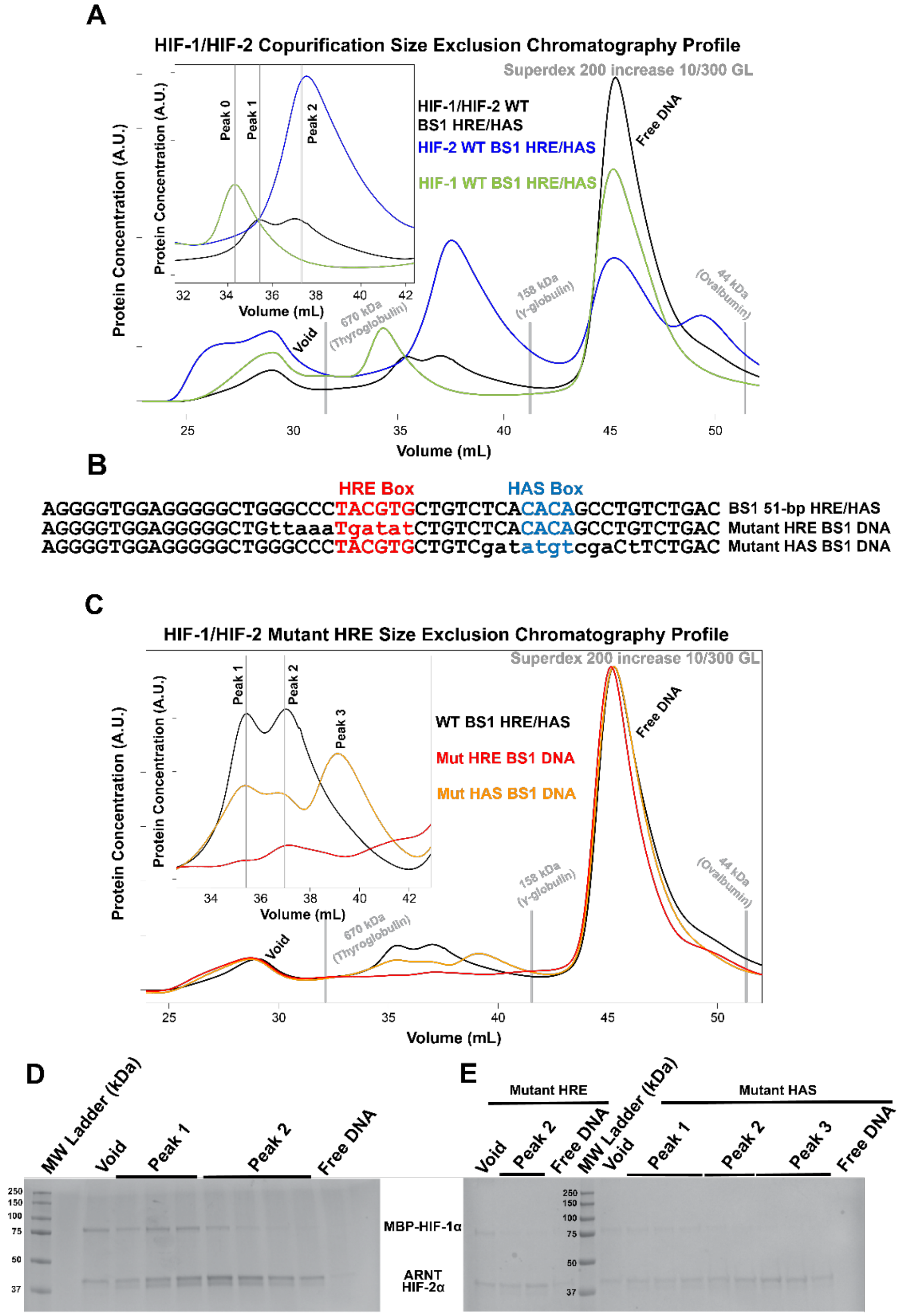


**Figure S9. Copurification of HIF-1 and HIF-2 on Extended DNA Fragments.** (A) Size exclusion chromatography profiles of copurified MBP-HIF-1 and HIF-2 (black), HIF-2 (blue), or MBP-HIF-1 (green) bound to 51-bp BS1 HRE/HAS. Peaks 0 and 2 correspond to a dimer of MBP-HIF-1 or HIF-2, respectively, while peak 1 is representative of a mixed MBP-HIF-1/HIF-2 dimer of heterodimers. (Expected complex sizes: MW MBP-HIF-1/MBP-HIF-1: 292.7 kDa, MW MBP-HIF-1/HIF-2: 247.7 kDa, MW HIF-2/HIF-2: 202.7 kDa) (B) Sequences of wildtype and mutant 51-bp BS1 HRE/HAS fragments. Differences from wildtype are represented as lowercase. (C) Size exclusion chromatography profiles of copurified MBP-HIF-1/HIF-2 incubated with wildtype 51-bp BS1 HRE/HAS (black), 51-bp BS1 DNA with a mutant HRE-box (red), or 51-bp BS1 DNA with a mutant HAS-box (orange). Peak 1 corresponds to the MBP-HIF-1/HIF-2 dimer of heterodimers, peak 2 corresponds to a HIF-2 dimer-of-heterodimers and we believe peak 3 corresponds to a single HIF heterodimer bound to DNA. (D) SDS-PAGE of fractions of the SEC of MBP-HIF-1/HIF-2 bound to wildtype 51-bp BS1 HRE/HAS (MBP-HIF-1α: 87.3 kDa, HIF-2α: 42.6 kDa, ARNT: 43.0 kDa). (E) SDS-PAGE of fractions of the SEC of MBP-HIF-1/HIF-2 bound to either mutant HRE-box (left) or mutant HAS-box (right) 51-bp BS1 DNA.

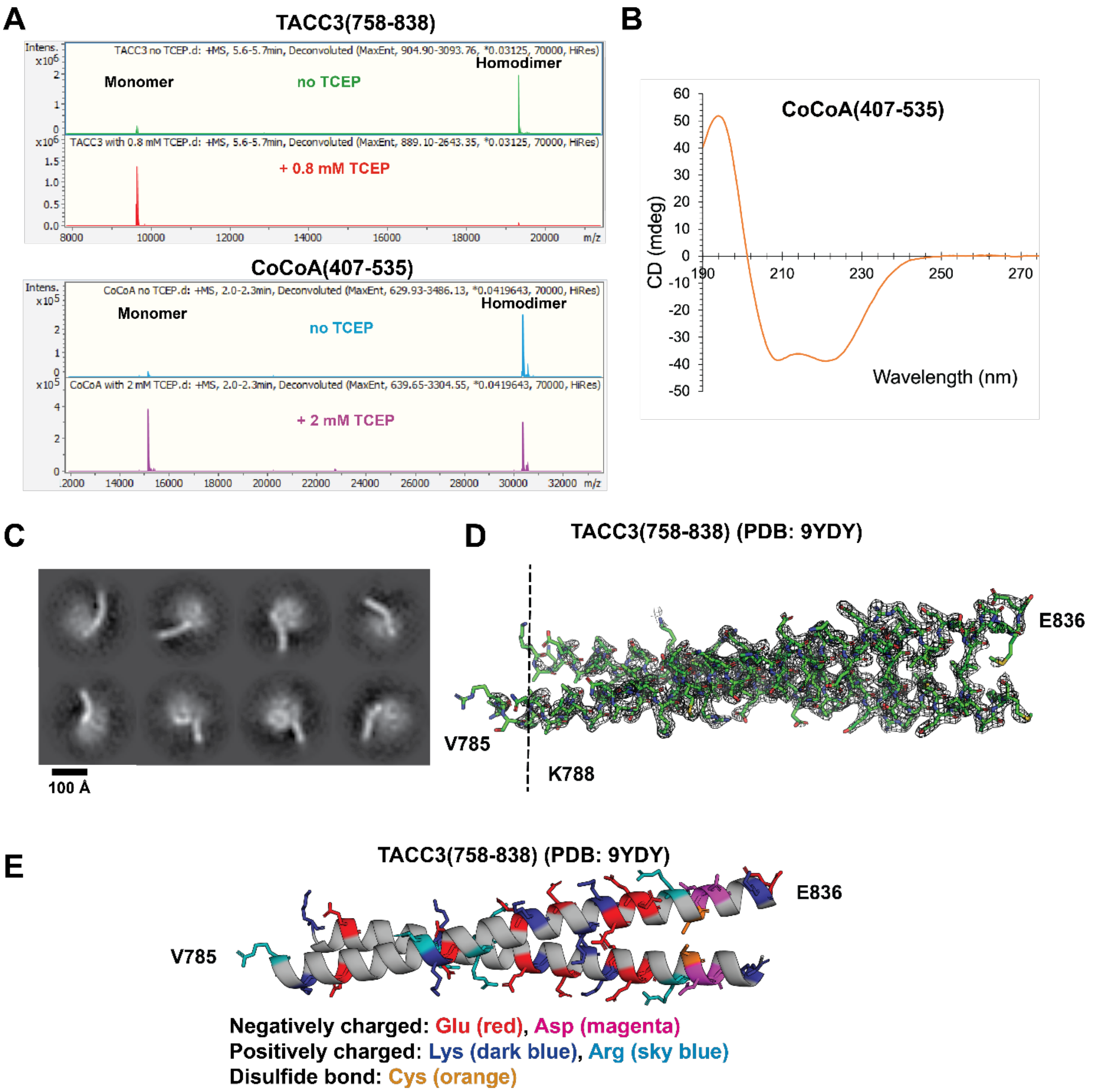


**Figure S10. Characterization of CCCs and CCC/HIF Complexes.** (A) Mass spectra of TACC3 and CoCoA with or without TCEP reducing agent. Reducing agent monomerizes both CCCs, but is more potent with TACC3. (B) Circular dichroism spectrum of CoCoA showing high presence of alpha helical structure, as expected from coiled coil structure. (C) Cryo-EM 2D classes of long TACC3(758-838) fragments bound to HIF-2 exhibit apparent bent stick-like TACC3(758-838) fragments bound to very blurred core domains, likely corresponding to HIF-2. Scale bar represents 100 Å. (D) X-ray density of TACC3(758-838) (PDB 9YDY) shows a loss of density N-terminal of lysine 788, suggesting that this region is dynamically unstable. (E) Mapping of charged residues visible within the TACC3(758-838) crystal structure.

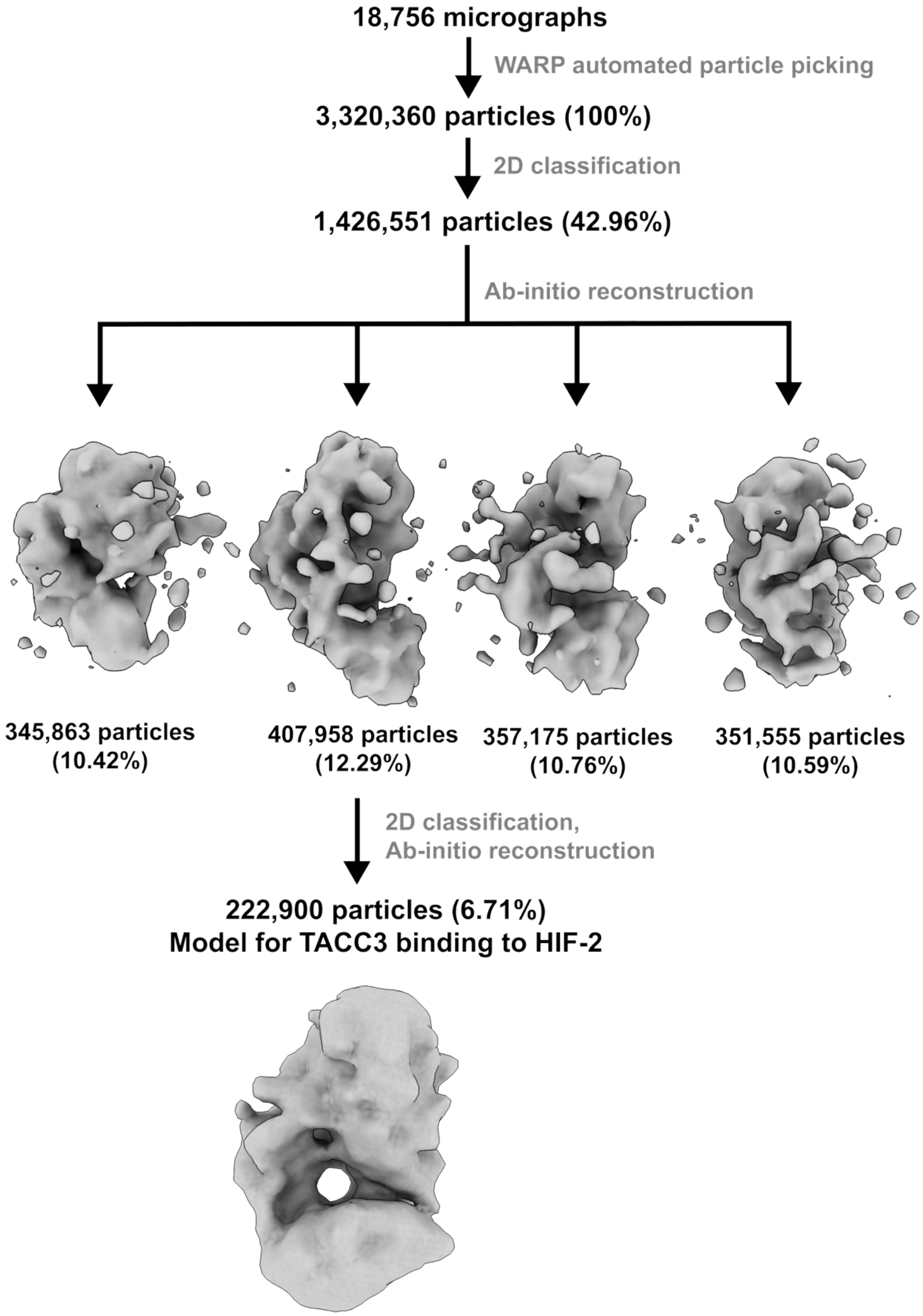


**Figure S11. HIF-2/20-bp DNA/TACC3 Cryo-EM Processing Workflow.** Particles picked from micrographs using WARP (5) were sorted into 3D classes using 2D classification and ab-initio reconstruction via CryoSPARC (6). Classes that appeared to contain excess density were further refined through several more rounds of 2D classification and ab-initio reconstruction, resulting in a cryo-EM map containing 222,900 particles that served as a model for TACC3 binding to HIF-2.

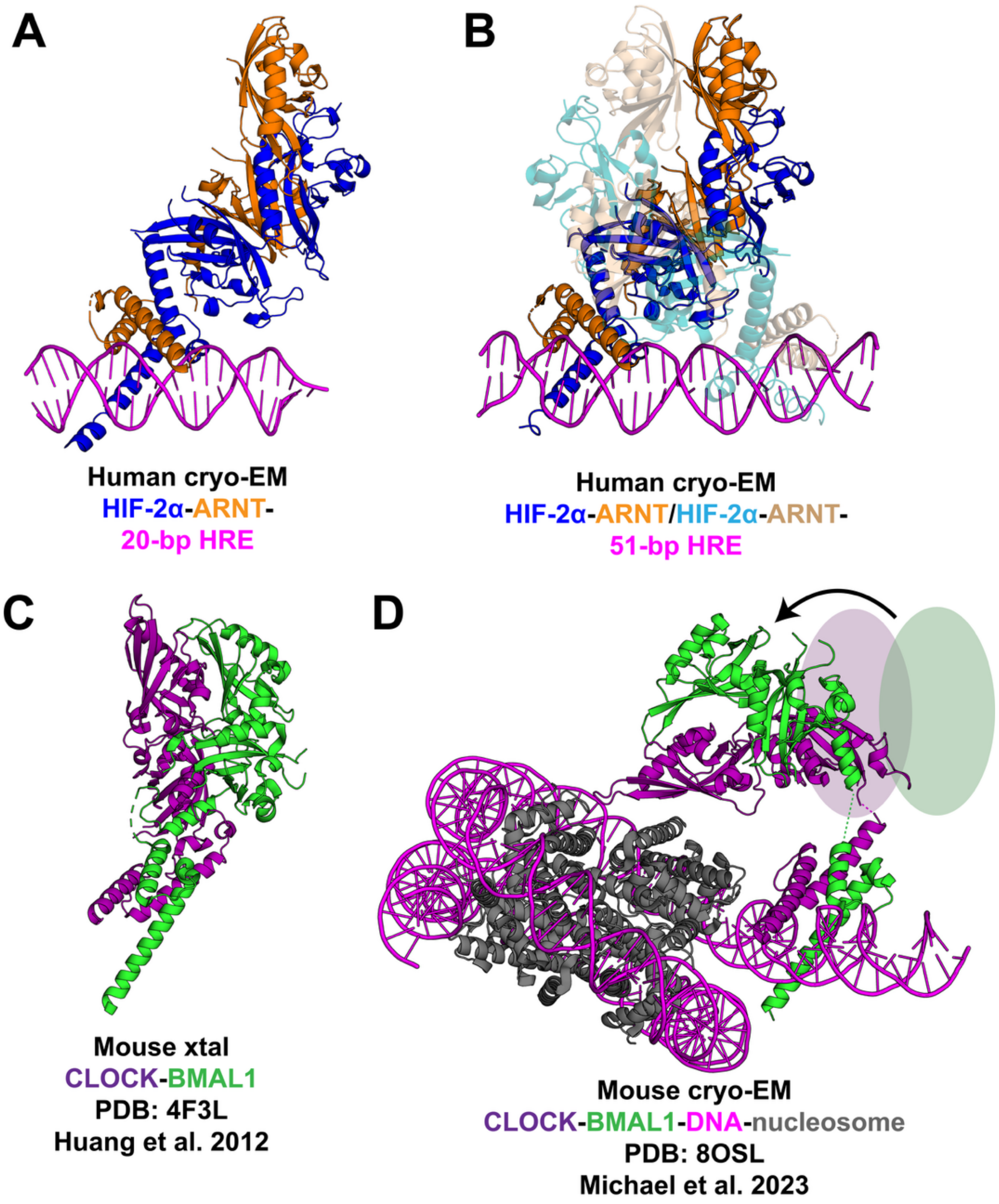


**Figure S12. Different bHLH-PAS Heterodimers Exhibit Different Levels of Conformational Change When Entering Higher Order Assemblies.** (A,B) Our cryo-EM structures of the human HIF-2 heterodimer (A) and dimer of heterodimers (B) show comparable arrangements of bHLH and PAS domains, suggesting a stable quaternary structure of the heterodimer that is effectively retained in an inverted orientation in the larger complex. (C,D). In contrast, the CLOCK-BMAL1 crystal structure solved in the absence of DNA (C) shows a marked rearrangement of the PAS domains with respect to the commonly-aligned bHLH domains when bound adjacent to a nucleosome (D) (4). Pale circles and arrow schematically represent required reorientation of PAS domains to convert from the DNA-free heterodimer to the nucleosome-bound one (8, 9).

| HRE-only Containing Genes | HRE Site Sequence | Function |
|---|---|---|
| GAPDH | TGCGTGCCCAGTTGAACCA | Glucose metabolism |
| TPI1 | CGCGTGGGTGTAGCTTTGT | Glucose metabolism |
| GPI | AACGTGGAAGGTGTTCGAT | Glucose metabolism |
| PGK1 | GACGTGACAAACGGAAGCC | Glucose metabolism |
| ENO1 | AGCGTGGAAAGGGGGGTGA | Glucose metabolism |
| PKM | TGCGTGGTTGGAGGTCCTG | Glucose metabolism |
| HK2 | AACGTGGGTATTTCCTGCT | Glucose metabolism |
| SLC16A4 | GGCGTGCGCCACTGTGCCC | Glucose metabolism |
| ALDOA | TGCGTGTCCTCTCGATGCC | Glucose metabolism |
| ALDOC | TACGTGACTCCTCCGGGGG | Glucose metabolism |
| PEPCK | GGCGTGTTGCAGTGAGTTG | Glucose metabolism |
| COX4I2 | GGCGTGAGCCACTGCTGGC | Glucose metabolism |
| CCNG2 | CACGTGCCCCGCCCCCTTG | Cell proliferation |
| TGFB1 | CGCGTGGGCGGGCTCCGAG | Cell proliferation |
| CDKN1A | AGCGTGCCCGCGTGTGTCC | Cell proliferation |
| TGFB2 | CACGTGGTTCAGAGAGAGA | Cell proliferation |
| ID2 | AGCGTGTGGCCCCGGCGAG | Cell proliferation |
| GADD45A (internal) | AGCGTGCCCCCCAACCCGA | Cell proliferation |
| CTGF | CGCGTGACTCAGGATGCAG | Cell proliferation |
| WT1 | AGCGTGTTTCAGAGGTGCG | Cell proliferation |
| IGFBP1 | AGCGTGCGGCGCTGCCAAT | Cell proliferation |
| IGFBP3 | GGCGTGCAGCTCGAGACTC | Cell proliferation |
| IGFBP4 | GGCGTGAGCCACCGCGCCC | Cell proliferation |
| ADM | CGCGTGGCTGAGGAAAGAA | Angiogenesis and $O_2$ transport |
| NOS2 | GGCGTGGTGGTTCATGCCG | Angiogenesis and $O_2$ transport |
| LRP1 | GGCGTGTCCCCTCGGGTGC | Angiogenesis and $O_2$ transport |
| TF | GACGTGTCCACTAGGCTTG | Angiogenesis and $O_2$ transport |
| EDN1 | GGCGTGGAGTGGGGTGGGA | Angiogenesis and $O_2$ transport |
| FECH | TGCGTGAAAATGGGCCCGG | Angiogenesis and $O_2$ transport |
| KDR | CACGTGTGTGGACTTCTTC | Angiogenesis and $O_2$ transport |
| FLT1 | CGCGTGGGGAGGTGCTGGC | Angiogenesis and $O_2$ transport |
| CYBB | GGCGTGAGCCCACCGTGCC | Angiogenesis and $O_2$ transport |
| GPX3 (downstream) | GACGTGGCTGGCATGGCCT | Angiogenesis and $O_2$ transport |
| AMFR | CACGTGCTAAGTCAAAGAA | Metastasis |
| ANGPTL4 | AGCGTGCGCCCTCTCCGCC | Metastasis |
| LGALS1 | AACGTGGTGAAACCCCGTC | Metastasis |
| LOX | AACGTGCAAGGCGGACAGC | Metastasis |
| LOXL4 | AACGTGGGCTCTGCCTGCT | Metastasis |
| MMP1 | AGCGTGACTCTGGGGTCTT | Metastasis |
| TWIST1 | GACGTGAGGAGGAGGGACT | Metastasis |
| COL5A1 | CACGTGTGGGTGAAGAGTG | Cellular structure |
| CD99 | CGCGTGGAGGCCGGGGCGG | Cellular structure |
| MMP2 | GACGTGGACATGAGCCCAG | Cellular structure |
| SERPINE1 | AACGTGAGCTGTTTTTTTT | Cellular structure |
| PLAUR | TGCGTGGGCGCCTCCCCCT | Cellular structure |
| KRT14 | GGCGTGATTCACAGGGGGA | Cellular structure |
| DELEC1 | GGCGTGAGCCACTGCGCCC | Transcriptional regulation |
| BHLHE41 | CACGTGCGGAACGTACCAT | Transcriptional regulation |
| CITED2 | CACGTGGCTGCAGAGGCTG | Transcriptional regulation |
| ETS1 | TGCGTGGGGCAAGGGGTTC | Transcriptional regulation |
| NR4A1 | CGCGTGCGTCACGCGCGCA | Transcriptional regulation |
| PMAIP1 | GGCGTGGCCTTACCGGGGA | Apoptosis |
| MCL1 | CACGTGCTACCCTAAAGAA | Apoptosis |
| BNIPL | GGCGTGTTGACCCAGCAGA | Apoptosis |
| NPPA | CACGTGGGCAGACCTCTGG | Cardiac Signaling |
| SNAI2 | TGCGTGCGGCGAGGGCGCT | Cell motility |
| PGP | TACGTGCCTGTTCAGTGAA | Drug resistance |
| TFF1 | AACGTGGTTCACTCCCCTG | Epithelial homeostasis |
| CXCL12 (downstream) | CGCGTGGAAAGCTGCGGGA | Inflammation |
| NT5E (downstream) | CACGTGGGGATTATGGGAG | Nucleotide metabolism |
| CA9 | AACGTGACCTAGTATGAGA | pH regulation |
| ANKRD37 | CACGTGCCAGTGTTTGTGT | Signaling |

| HRE/HAS Containing Genes | HRE/HAS Site Sequence | Function |
|---|---|---|
| PFKL (internal) | TACGTGCGGCCGGCCACGT | Glucose metabolism |
| HK1 (downstream) | GGCGTGCATCCCCACACCC | Glucose metabolism |
| LDHA | CACGTGGGTTCCCGCACGT | Glucose metabolism |
| PGM1 (internal) | GGCGTGAGCCACCACACAC | Glucose metabolism |
| SLC2A3 (internal) | GGCGTGAGCCACGGCACCG | Glucose metabolism |
| SLC9A1 | GGCGTGGTGGCAGGCACCT | Glucose metabolism |
| PFKFB1 (downstream) | GACGTGCCTGACTCCACAC | Glucose metabolism |
| PGM (downstream) | GGCGTGTGCCACCACACCT | Glucose metabolism |
| TKT (internal) | AACGTGACCCACGCCACAA | Glucose metabolism |
| TKTL2 | GGCGTGGTGATGGGCACCT | Glucose metabolism |
| MXI1 (downstream) | GGCGTGGTGGCATGCACCT | Glucose metabolism |
| LONP (internal) | GGCGTGGTGGCGGGCACCT | Glucose metabolism |
| SLC20A1 (internal) | CACGTGCACACACACACTC | Glucose metabolism |
| GLUT1/SLC2A1 | GGCGTGGTGGCTCTCACCT | Glucose metabolism |
| EPO (downstream) | TACGTGCTGTCTCACACAG | Angiogenesis and $O_2$ transport |
| VEGFB (internal) | GGCGTGGTGGCGGGCACCT | Angiogenesis and $O_2$ transport |
| CP (downstream) | GACGTGGTGGTGCGCACCT | Angiogenesis and $O_2$ transport |
| HMOX1 (internal) | GACGTGGCACTTATCACAG | Angiogenesis and $O_2$ transport |
| TFRC (downstream) | GGCGTGGTGGCAGACACCT | Angiogenesis and $O_2$ transport |
| ANGPT1 | CGCGTGCGCGCGCACACGC | Angiogenesis and $O_2$ transport |
| NOS3 (internal) | GACGTGAGCCACTGCACCT | Angiogenesis and $O_2$ transport |
| PDGFB (internal) | CACGTGTGCAAACACACAC | Angiogenesis and $O_2$ transport |
| PGF | CAGGTGCAGTGGCTCACGC | Angiogenesis and $O_2$ transport |
| SOD2 | GGCGTGGTGGCACGCACCT | Angiogenesis and $O_2$ transport |
| VEGFC (internal) | GGCGTGGTGGTTCACACTG | Angiogenesis and $O_2$ transport |
| ANGPT2 (internal) | GGCGTGATCCACCACACCT | Angiogenesis and $O_2$ transport |
| IGFBP2 | GGCGTGCGCGCACTCACTT | Cell proliferation |
| TGFA (downstream) | GGCGTGGTGGCGCACACCT | Cell proliferation |
| TGFB3 | CACGTGGCTCCCCGCACCC | Cell proliferation |
| ABCG2 (downstream) | GGCGTGGTGGCACACACT | Cell proliferation |
| POU5F1 | GGCGTGAGCCACCACACCC | Cell proliferation |
| TERT | GACGTGGAAAGGAGCACAG | Cell proliferation |
| DDIT4 | GACGTGGTGGTGGGCACCT | Cell proliferation |
| RORA | GGCGTGAGCCACCGCACCT | Cell proliferation |
| AURKA (internal) | AACGTGAACTCCAACACAG | Cell proliferation |
| CTSC (internal) | GGCGTGATGGCGGGCACCT | Metastasis |
| CXCR4 | CGCGTGCGTCGCGACACGC | Metastasis |
| L1CAM (internal) | GACGTGTTGGACACCACCA | Metastasis |
| LOXL2 | GACGTGAGCCACTTCACCT | Metastasis |
| STC2 | TACGTGTGCACGCTCACGT | Metastasis |
| MET (downstream) | GGCGTGAGCCACCACACCC | Cell motility |
| SNAI1 (internal) | AGCGTGAGCCACCACACCC | Cell motility |
| TCF3 | GGCGTGGTGGCATACACCT | Cell motility |
| ZEB1 (downstream) | GGCGTGGTGGCAGGCACCT | Cell motility |
| ZEB2 (internal) | GGCGTGGTTGCAGGCACCT | Cell motility |
| VIM (downstream) | GGCGTGGTGGCTCACACTT | Cellular structure |
| P4HA1 | TACGTGAGCTCGATCACAC | Cellular structure |
| PLAUR | GGCGTGAGCCACTGCACCC | Cellular structure |
| CTSD | GGCGTGAGTAATCCCACTT | Cellular structure |
| BNIP3 (internal) | AGCGTGCATATAGGCACAC | Apoptosis |
| NDRG1 (downstream) | GGCGTGAGCCACCGCACCC | Apoptosis |
| PPP5C (internal) | AGCGTGGTGGCACGCACCT | Apoptosis |
| NPM | GGCGTGGTGGCATGCACCT | Anti-apoptosis |
| LEP (internal) | GGCGTGATGACTCACACCT | Fatty acid metabolism |

**Table S1. Characterization of HRE-box or HRE/HAS Motif Containing HIF Genes.** *In-silico* survey of HIF-1 or HIF-2 controlled human genes, conducted to determine if the promoter (upstream) or enhancer (downstream/internal) regions contained only an HRE-box (left, most proximal HRE-box chosen in case of multiples) or an HRE-box followed by an 8 bp spacer and HAS-box (right). Several displayed genes (yellow highlights) have been experimentally validated as being HIF regulated (EPO (10), PFKL (11), HK1 (12), VEGF (13), CA9 (12), PGK1 (14), BNIP3 (15)). All genes were sorted by cellular function.

| | **HIF-2/HRE**<br>PDB 9OF0<br>EMD-70416 | **HIF-2/HRE+HAS**<br>PDB 9OF2<br>EMD-70418 | **HIF-1/HRE+HAS**<br>PDB 9OFU<br>EMD-70443 |
|---|---|---|---|
| **Data collection and processing** | | | |
| Magnification | 105,000x | 105,000x | 105,000x |
| Voltage (kV) | 300 | 300 | 300 |
| Electron exposure ($e^{-}/Å^{2}$) | 55 | 59 | 50 |
| Defocus range (μm) | 0 to -2.0 | 0 to -2.0 | 0 to -2.0 |
| Pixel size (Å) | 0.844 | 0.83 | 0.829 |
| Symmetry imposed | C1 | C1 | C1 |
| Initial micrographs | 13,373 | 10,910 | 4,098 |
| Final micrographs | 13,373 | 10,910 | 4,098 |
| Initial particle images | 1.98M | 657K | 253K |
| Final particle images | 26K | 164K | 72K |
| Subparticle images | NA | NA | NA |
| Map resolution (Å) | 3.6 | 3.7 | 3.9 |
| **Model composition** | | | |
| Non-hydrogen atoms | 5,572 | 10,149 | 10,067 |
| Protein residues | 582 | 1,115 | 1,067 |
| **RMS Deviations** | | | |
| Bond lengths (Å) | 0.21 | 0.19 | 0.14 |
| Bond angles (º) | 0.38 | 0.34 | 0.31 |
| **Validation** | | | |
| Clash score | 7 | 8 | 9 |
| Poor rotamers (%) | 1.3 | 1.2 | 1 |
| **Ramachandran plot** | | | |
| Favored (%) | 98 | 96 | 98 |
| Allowed (%) | 2 | 4 | 2 |
| Disallowed (%) | 0 | 0 | 0 |

**Table S2. Collection, Processing and Refinement Statistics for Cryo-EM Structures.** NA = not applicable.

| | **TACC3(758-838)** |
|---|---|
| Wavelength (Å) | 0.9201 |
| Space Group | C2 |
| Unit cell a, b ,c (Å) | 263.08, 23.83, 31.86 |
| Unit cell $\alpha$, $\beta$, $\gamma$ (°) | 90, 91.81, 90 |
| Resolution range (Å) | 32.87 – 2.30 (2.38 – 2.30) |
| Total reflections | 30,861 |
| Unique reflections | 9,352 |
| Mean(I)/$\sigma$(I) | 7.00 (1.20) |
| Completeness (%) | 98.4 (93) |
| Multiplicity | 3.3 |
| $CC_{1/2}$ | 0.992 (0.580) |
| $R_{merge}$ | 0.085 |
| Reflections used in refinement | 9011 |
| Reflections used for $R_{free}$ | 911 |
| $R_{work}$ | 0.254 |
| $R_{free}$ | 0.292 |
| Number of nonhydrogen atoms | 819 |
| Macromolecules | 802 |
| Solvent | 17 |
| Protein residues | 100 |
| RMS (bonds) | 0.009 |
| RMS (angles) | 1.15 |
| Ramachandran favored (%) | 100 |
| Ramachandran allowed (%) | 0 |
| Rotamer outliers (%) | 0 |
| Clash score | 4.86 |
| Average B-factor (Å$^2$) | 67.24 |
| Macromolecules | 67.45 |
| Solvent | 57.17 |
| PDB accession code | 9OPF |

**Table S3. Crystallographic data collection, processing, and refinement statistics for structure of TACC3(758-838).** Structure includes residues 788-836 of chain A, and residues 785-836 of chain B.

| Protein | Residues | Deuteration Time (sec) | #D (no TACC3) | %D (no TACC3) | #D (w/TACC3) | %D (w/TACC3) |
|---|---|---|---|---|---|---|
| ARNT | 330-341 GQGSKFCLVAIG | 0 | 0 (+/- 0) | 0 | 0 (+/- 0) | 0 |
| | | 50 | 4.856 (+/- 0.238) | 48.559 | 2.54 (+/- 0.066) | 33.418 |
| | | 300 | 5.003 (+/- 0.4) | 50.034 | 2.668 (+/- 0.075) | 35.103 |
| | | 1000 | 5.237 (+/- 0.196) | 52.367 | 2.669 (+/- 0.087) | 35.122 |
| | | 3000 | 5.499 (+/- 0.34) | 54.992 | 2.511 (+/- 0.128) | 33.041 |
| | | 5000 | 5.839 (+/- 0.313) | 58.386 | 2.699 (+/- 0.116) | 35.512 |
| ARNT | 357-366 VCQPTEFISR | 0 | 0 (+/- 0) | 0 | 0 (+/- 0) | 0 |
| | | 50 | 3.919 (+/- 0.506) | 58.926 | 2.949 (+/- 0.056) | 44.352 |
| | | 300 | 3.977 (+/- 0.294) | 59.806 | 3.331 (+/- 0.123) | 50.088 |
| | | 1000 | 5.149 (+/- 0.444) | 77.436 | 3.934 (+/- 0.12) | 59.158 |
| | | 3000 | 5.458 (+/- 0.366) | 82.083 | 4.55 (+/- 0.225) | 68.425 |
| | | 5000 | 5.283 (+/- 0.26) | 79.437 | 4.873 (+/- 0.065) | 73.278 |
| ARNT | 372-379 IFTFVDHR | 0 | 0 (+/- 0) | 0 | 0 (+/- 0) | 0 |
| | | 50 | 1.125 (+/- 0.125) | 19.73 | 0.665 (+/- 0.035) | 11.659 |
| | | 300 | 1.156 (+/- 0.299) | 20.286 | 0.788 (+/- 0.04) | 13.833 |
| | | 1000 | 1.201 (+/- 0.235) | 21.077 | 0.981 (+/- 0.006) | 17.216 |
| | | 3000 | 1.671 (+/- 0.375) | 29.318 | 1.346 (+/- 0.021) | 23.622 |
| | | 5000 | 1.751 (+/- 0.114) | 30.718 | 1.52 (+/- 0.014) | 26.669 |
| ARNT (control) | 287-294 NRCRNGLG | 0 | 0 (+/- 0) | 0 | 0 (+/- 0) | 0 |
| | | 50 | 0.639 (+/- 0.169) | 10.648 | 0.568 (+/- 0.091) | 9.962 |
| | | 300 | 0.62 (+/- 0.146) | 10.342 | 0.561 (+/- 0.048) | 9.841 |
| | | 1000 | 0.714 (+/- 0.139) | 11.894 | 0.668 (+/- 0.07) | 11.728 |
| | | 3000 | 0.848 (+/- 0.159) | 14.126 | 0.816 (+/- 0.038) | 14.317 |
| | | 5000 | 0.962 (+/- 0.1) | 16.036 | 0.96 (+/- 0.063) | 16.85 |
| ARNT (control) | 205-225 FGSTLYDQVHPDD VDKLREQL | 0 | 0 (+/- 0) | 0 | 0 (+/- 0) | 0 |
| | | 50 | 2.599 (+/- 0.347) | 14.436 | 2.496 (+/- 0.108) | 14.596 |
| | | 300 | 3.319 (+/- 0.287) | 18.439 | 3.411 (+/- 0.093) | 19.95 |
| | | 1000 | 3.781 (+/- 0.312) | 21.008 | 3.953 (+/- 0.136) | 23.117 |
| | | 3000 | 4.478 (+/- 0.46) | 24.878 | 4.724 (+/- 0.17) | 27.628 |
| | | 5000 | 5.012 (+/- 0.446) | 27.845 | 5.3 (+/- 0.191) | 30.996 |
| ARNT (control) | 150-167 TDGSYKPSFLTDQE LKHL | 0 | 0 (+/- 0) | 0 | 0 (+/- 0) | 0 |
| | | 50 | 7.019 (+/- 0.773) | 46.791 | 6.774 (+/- 0.554) | 47.536 |
| | | 300 | 6.969 (+/- 0.789) | 46.458 | 6.969 (+/- 0.394) | 48.908 |
| | | 1000 | 7.159 (+/- 0.775) | 47.729 | 7.133 (+/- 0.463) | 50.059 |
| | | 3000 | 7.109 (+/- 0.734) | 47.395 | 7.242 (+/- 0.373) | 50.82 |
| | | 5000 | 7.001 (+/- 0.883) | 46.673 | 7.474 (+/- 0.543) | 52.446 |
| HIF-2α | 155-162 FGKKSKDM | 0 | 0 (+/- 0) | 0 | 0 (+/- 0) | 0 |
| | | 50 | 1.754 (+/- 0.073) | 29.226 | 1.522 (+/- 0.029) | 26.703 |
| | | 300 | 2.112 (+/- 0.03) | 35.193 | 1.939 (+/- 0.006) | 34.024 |
| | | 1000 | 2.428 (+/- 0.036) | 40.466 | 2.276 (+/- 0.039) | 39.921 |
| | | 3000 | 2.668 (+/- 0.091) | 44.474 | 2.593 (+/- 0.034) | 45.484 |
| | | 5000 | 2.893 (+/- 0.056) | 48.217 | 2.848 (+/- 0.04) | 49.966 |
| HIF-2α | 157-163 KKSKDMS | 0 | 0 (+/- 0) | 0 | 0 (+/- 0) | 0 |
| | | 50 | 1.588 (+/- 0.196) | 33.438 | 1.23 (+/- 0.022) | 25.891 |
| | | 300 | 1.516 (+/- 0.039) | 31.918 | 1.255 (+/- 0.013) | 26.417 |
| | | 1000 | 1.464 (+/- 0.05) | 30.815 | 1.285 (+/- 0.04) | 27.06 |
| | | 3000 | 1.558 (+/- 0.059) | 32.799 | 1.312 (+/- 0.028) | 27.631 |
| | | 5000 | 1.597 (+/- 0.028) | 33.612 | 1.373 (+/- 0.045) | 28.9 |
| HIF-2α | 201-208 VYNNCPPH | 0 | 0 (+/- 0) | 0 | 0 (+/- 0) | 0 |
| | | 50 | 3.64 (+/- 0.206) | 90.998 | 2.327 (+/- 0.045) | 61.231 |
| | | 300 | 3.65 (+/- 0.228) | 91.258 | 2.664 (+/- 0.092) | 70.115 |
| | | 1000 | 3.575 (+/- 0.389) | 89.37 | 2.708 (+/- 0.162) | 71.261 |
| | | 3000 | 3.856 (+/- 0.32) | 96.402 | 2.92 (+/- 0.187) | 76.835 |
| | | 5000 | 3.91 (+/- 0.239) | 97.741 | 2.967 (+/- 0.281) | 78.082 |
| HIF-2α | 221-229 CLIIMCEPI | 0 | 0 (+/- 0) | 0 | 0 (+/- 0) | 0 |
| | | 50 | 0.538 (+/- 0.461) | 8.964 | -0.128 (+/- 0.069) | -2.253 |
| | | 300 | 0.39 (+/- 0.14) | 6.504 | 0.009 (+/- 0.048) | 0.163 |
| | | 1000 | 0.518 (+/- 0.216) | 8.634 | 0.178 (+/- 0.025) | 3.118 |
| | | 3000 | 1.807 (+/- 0.528) | 30.117 | 0.719 (+/- 0.1) | 12.615 |
| | | 5000 | 2.322 (+/- 0.521) | 38.7 | 0.789 (+/- 0.038) | 13.836 |
| HIF-2α | 323-333 GTVIYNPRNLQ | 0 | 0 (+/- 0) | 0 | 0 (+/- 0) | 0 |
| | | 50 | 5.268 (+/- 0.436) | 65.845 | 7.087 (+/- 0.148) | 93.251 |
| | | 300 | 5.228 (+/- 0.319) | 65.345 | 7.034 (+/- 0.178) | 92.558 |
| | | 1000 | 5.409 (+/- 0.328) | 67.616 | 7.145 (+/- 0.152) | 94.011 |

| | | | | | | |
|---|---|---|---|---|---|---|
| | | 3000 | 5.488 (+/- 0.424) | 68.599 | 7.16 (+/- 0.152) | 94.21 |
| | | 5000 | 5.625 (+/- 0.385) | 70.307 | 7.211 (+/- 0.117) | 94.88 |
| HIF-2α | 346-355 EIEKNDVVFS | 0 | 0 (+/- 0) | 0 | 0 (+/- 0) | 0 |
| | | 50 | 3.463 (+/- 1.082) | 43.293 | 2.719 (+/- 0.102) | 35.774 |
| | | 300 | 3.904 (+/- 0.244) | 48.8 | 3.269 (+/- 0.429) | 43.015 |
| | | 1000 | 4.624 (+/- 0.483) | 57.794 | 3.427 (+/- 0.153) | 45.098 |
| | | 3000 | 4.809 (+/- 0.265) | 60.108 | 4.02 (+/- 0.218) | 52.898 |
| | | 5000 | 4.886 (+/- 0.691) | 61.074 | 4.651 (+/- 0.193) | 61.192 |
| HIF-2α (control) | 300-309 GQVVSGQYRM | 0 | 0 (+/- 0) | 0 | 0 (+/- 0) | 0 |
| | | 50 | 3.582 (+/- 1.613) | 44.774 | 1.733 (+/- 0.145) | 22.796 |
| | | 300 | 2.112 (+/- 0.138) | 26.396 | 1.791 (+/- 0.331) | 23.561 |
| | | 1000 | 2.385 (+/- 0.141) | 29.819 | 2.315 (+/- 0.051) | 30.46 |
| | | 3000 | 2.832 (+/- 0.252) | 35.402 | 2.786 (+/- 0.12) | 36.66 |
| | | 5000 | 3.000 (+/- 0.157) | 37.495 | 2.981 (+/- 0.069) | 39.226 |
| HIF-2α (control) | 216-235 EPLLSCLIIMCEPIQ HPSHM | 0 | 0 (+/- 0) | 0 | 0 (+/- 0) | 0 |
| | | 50 | 3.749 (+/- 1.071) | 23.432 | 3.496 (+/- 0.222) | 23.003 |
| | | 300 | 4.474 (+/- 0.932) | 27.964 | 4.545 (+/- 0.152) | 29.901 |
| | | 1000 | 5.441 (+/- 1.071) | 34.008 | 5.488 (+/- 0.323) | 36.105 |
| | | 3000 | 6.447 (+/- 1.668) | 40.295 | 6.696 (+/- 0.208) | 44.056 |
| | | 5000 | 7.15 (+/- 1.392) | 44.686 | 7.718 (+/- 0.154) | 50.777 |
| HIF-2α (control) | 111-121 LSENISKFMGL | 0 | 0 (+/- 0) | 0 | 0 (+/- 0) | 0 |
| | | 50 | 2.309 (+/- 0.174) | 25.658 | 2.118 (+/- 0.222) | 24.775 |
| | | 300 | 2.363 (+/- 0.235) | 26.254 | 2.227 (+/- 0.095) | 26.05 |
| | | 1000 | 2.474 (+/- 0.218) | 27.488 | 2.415 (+/- 0.07) | 28.24 |
| | | 3000 | 2.481 (+/- 0.3) | 27.572 | 2.495 (+/- 0.106) | 29.182 |
| | | 5000 | 2.701 (+/- 0.224) | 30.015 | 2.682 (+/- 0.099) | 31.368 |

**Table S4. Selected Hydrogen-Deuterium Exchange Data for HIF-2 peptides in the absence and presence of TACC3.** Comparison of deuteration levels of select peptides of HIF-2α and ARNT in the absence (columns 4, 5) and presence (columns 6, 7) of TACC3. Absolute deuteration levels (columns 4, 6), are reported as averages and standard deviation of number of incorporated deuterons from three independent replicates; these are also reported as % average deuteration in columns 5, 7. Control peptides are included in which there is minimal change in deuteration upon adding TACC3.
